# Integrating stool chemistry measurements with metagenomics in adults reveals distinct gut health profiles

**DOI:** 10.64898/2026.10.23.759375

**Authors:** Timothy J. Straub, Pamela A. Nieto, Julian M. Trachsel, Hannah Sturgeon, Star Edwards, Emeran A. Mayer, Ruben A. T. Mars, Cheryl Sew Hoy, Kimberley V. Sukhum

**Affiliations:** Seeding Inc, doing business as Tiny Health, Austin, Texas, USA; G. Oppenheimer Center for Neurobiology of Stress and Resilience, UCLA, Los Angeles, CA, USA; Goodman-Luskin Microbiome Center, UCLA, Los Angeles, CA, USA; Vatche and Tamar Manoukian Division of Digestive Diseases, UCLA, Los Angeles, CA, USA; David Geffen School of Medicine, UCLA, Los Angeles, CA, USA; University of California, Los Angeles, CA, USA; Division of Gastroenterology and Hepatology, Mayo Clinic, Rochester, Minnesota, USA

## Abstract

Stool chemistry metrics, including calprotectin, lactoferrin, and secretory IgA, are used to assess gut inflammation in clinical or research settings. However, these measures in isolation provide a limited view of gut physiology and lack microbial context. Shotgun metagenomics offers a comprehensive assessment of the gut microbiome’s taxonomic composition and functional potential, yet its integration with stool chemistry metrics remains underexplored. We analyzed 877 adult stool samples with paired shotgun metagenomic sequencing and chemistry measurements to integrate the microbiome with 12 stool chemistry metrics. Calprotectin, lactoferrin, and secretory IgA were associated with shifts in microbiome composition (beta diversity), while deviations from typical levels of short-chain fatty acids (SCFAs), including butyrate, were associated with both changes in alpha diversity and microbiome composition. Network analysis revealed two distinct microbial neighborhoods: one associated with higher butyrate and lower calprotectin and lactoferrin and another associated with lower butyrate and higher calprotectin and lactoferrin. Machine learning models trained on taxonomic and functional data showed modest discrimination of stool chemistry states, performing best for SCFA deficiency and less well for host-derived inflammatory metrics. Together, these findings systematically characterize microbiome-stool chemistry relationships across a large paired dataset, providing a framework for interpreting stool chemistry metrics in the context of shotgun metagenomic profiles.

## Introduction

The human gut microbiome is fundamental to health, contributing to digestion, immune regulation, and metabolic balance^1^. Alterations in microbial composition and function have been associated with a wide range of conditions, including inflammatory bowel disease (IBD), irritable bowel syndrome (IBS)^2^, and metabolic disorders such as type 2 diabetes and obesity^3^. Despite these associations, clinical evaluation of gut health often remains limited to symptom-based assessment or measurement of a few stool chemistry metrics, approaches that may overlook microbial contributions to disease and lead to ambiguous interpretations.

Among stool chemistry metrics used in clinical practice, calprotectin is a well-established biomarker of intestinal inflammation, routinely used to distinguish IBD from functional disorders such as IBS, and to monitor disease activity^4–6^. Stool lactoferrin, another neutrophil-derived protein, similarly reflects mucosal inflammation and has been proposed as a supportive inflammatory marker, although it is less commonly used in routine clinical workflows^5,7^. Together, these host-derived proteins serve as sensitive indicators of intestinal immune activation but do not provide information about which specific microbial taxa or functional features may be contributing to inflammatory states. Additional stool chemistry metrics—such as lysozyme, secretory IgA (sIgA), and stool pH—are sometimes used in research to capture aspects of mucosal immunity, barrier function, and microbial metabolism; however, their clinical interpretation and mechanistic links to underlying microbiome features remain less well defined.

In contrast, short-chain fatty acids (SCFAs)—including butyrate, propionate, acetate, and valerate—are stool chemistry metrics that represent microbial metabolic outputs. Mostly produced by bacterial fermentation of dietary fibers, SCFAs represent a key link between the microbiome and host physiology: they fuel colonocytes, reinforce the intestinal barrier, and modulate immune responses^8^. That said, stool SCFA profiles can be highly variable and influenced by both host and microbial factors, raising questions about how well they reflect *in vivo* production and activity^9^.

These distinctions highlight a key challenge in gut health assessment: host-derived inflammatory metrics and microbial metabolic readouts capture different biological layers yet are rarely interpreted together or linked to underlying microbial mechanisms. Shotgun metagenomic sequencing offers a way to bridge this gap by providing species- and strain-level taxonomic resolution alongside the functional genetic potential of the entire microbial community^10^. Unlike targeted assays or amplicon-based approaches (e.g., qPCR or 16S rRNA), metagenomics enables direct interrogation of microbial taxa and pathways that may be associated with host inflammatory responses or influence metabolite availability. Large multi-omics studies, including disease-focused cohorts such as the Integrative Human Microbiome Project (HMP2), have demonstrated the value of integrating microbiome data with host-derived measurements^11,12^. However, few studies have systematically examined a broad panel of clinically relevant stool chemistry metrics alongside shotgun metagenomics across a heterogeneous adult population. As a result, clinicians and researchers often interpret stool chemistry metrics and metagenomic profiles independently, limiting their ability to place biomarker abnormalities into a broader microbial and functional context.

To address this gap, we performed an exploratory characterization of 877 adult stool samples with paired stool chemistry measurements and shotgun metagenomic profiles. Our goal was to characterize associations between stool chemistry metrics and the gut microbiome, including microbial diversity, taxonomic composition, and functional potential. As a secondary objective, we evaluated whether metagenomic features could predict selected stool chemistry metrics using machine learning approaches. Together, these analyses provide a descriptive overview of how stool chemistry measurements relate to gut microbial structure and function in a large paired metagenomics – stool chemistry cohort.

## Results

### Study population, health conditions, and demographics

To investigate the relationships between metagenomics and stool chemistry metrics, we obtained stool samples from 877 adult participants through convenience sampling. Demographic, lifestyle, and self-reported health data were collected for all individuals (**Fig. 1a**; **Table 1; Suppl. File 1, Table S1**). The mean age was 41.3 years (SD = 11.3; range, 18–64), and the cohort was composed primarily of females (76.2%, n = 668). The average Body Mass Index (BMI) for the cohort was 25.2 kg/m² (SD = 5.8). The cohort was predominantly White (73%, with other represented ethnicities including Hispanic or Latino, 9.5%; Asian, 6.7%; Black or African American, 1.3%; and other or unknown identities) (**Table 1**). Participants self-reported a variety of health conditions, which were grouped into predefined categories (**Suppl. File 1, Table S2**). Because participants could report multiple conditions spanning different categories, category counts are not mutually exclusive (**Fig. 1b**). The most frequently reported categories were general digestive issues (62%), allergies or atopic conditions (45.7%), and gynecological and hormonal conditions (28.5%). A diagnosis of IBD was reported by 6.6% (n = 58) of participants. Notably, 10.3% (n = 90) of the cohort self-identified as healthy (**Table 1**). While 40% of the healthy group reported taking at least one medication, medication use exceeded 70% among participants with metabolic, mental/neurological, and other inflammatory/autoimmune conditions. Use of proton pump inhibitor (PPI)/reflux medications and NSAIDs, known to influence calprotectin levels and gut microbiome composition^13–15^, was highest among participants who reported metabolic conditions (22%) **(Suppl. File 1, Table S1**).

**Fig. 1:**
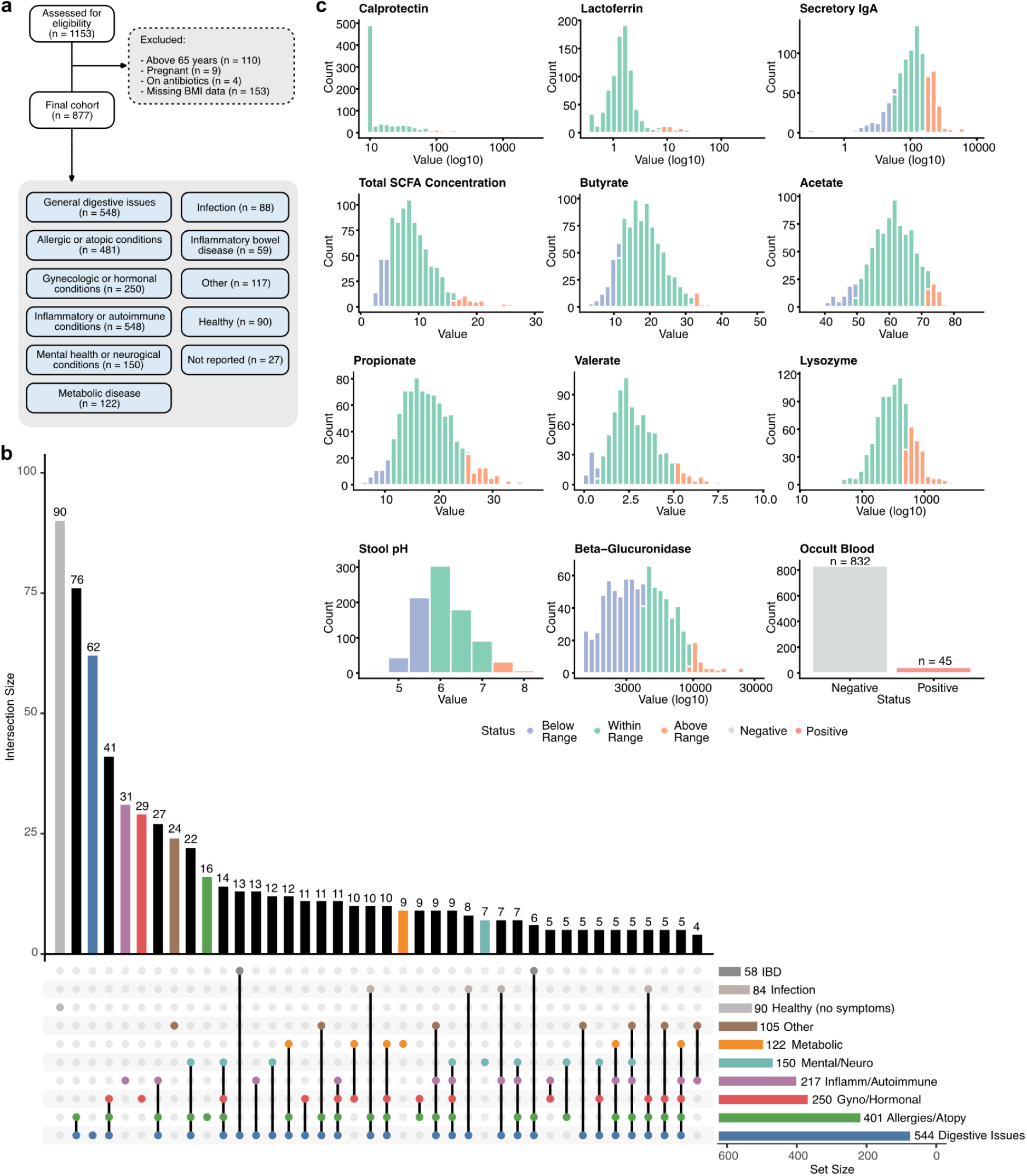
Study cohort shows substantial heterogeneity in self-reported health conditions but largely normal stool chemistry. **(a)** Participant selection flow diagram showing eligibility assessment, exclusion criteria, and final cohort size (n = 877). Self-reported health conditions were grouped into predefined categories; participants could report multiple conditions, and categories are therefore not mutually exclusive. (b) Upset plot showing the distribution and overlap of self-reported health condition categories across the cohort. Bar heights indicate intersection sizes, and horizontal bars show the total number of participants reporting each category. (c) Distributions of stool chemistry metrics across all samples. Continuous variables are shown on either a linear or log₁₀ scale, as indicated, to accommodate skewed distributions. Binary measures (e.g., positive vs negative) are shown as count distributions by category. Samples were grouped as below range (blue), within range (green), or above range (orange) based on reference ranges for each chemistry metric.

**Table 1.** Baseline demographic and self-reported clinical characteristics of the study cohort.

| Characteristic | Overall<br>(n=877, 100%) | Healthy<br>(n=90, 10.2%) | General<br>Digestive<br>Issues<br>(n=544, 62%) | Allergies/Atopic<br>(n=401, 45.7%) | Gyno/<br>Hormonal<br>(n=250, 28.5%) | Other<br>Inflam/<br>Autoimmune<br>(n=217, 24.7%) | Mental<br>/Neuro<br>(n=150, 17.1%) | Metabolic<br>(n=122, 13.9%) | Infectious<br>(n=84, 9.6%) | IBD<br>(n=58, 6.6%) | Other<br>(n=105, 12%) |
| --- | --- | --- | --- | --- | --- | --- | --- | --- | --- | --- | --- |
| Age, years | 41.3 (11.3) | 40.7 (10.8) | 41.4 (11.2) | 41.1 (11.1) | 38.9 (10.0) | 43.8 (10.2) | 37.0 (10.7) | 42.4 (11.0) | 42.1 (10.9) | 40.4 (11.6) | 41.7 (11.8) |
| Sex |  |  |  |  |  |  |  |  |  |  |  |
| Female | 668 (76%) | 60 (67%) | 427 (78%) | 327 (82%) | 250 (100%) | 184 (85%) | 114 (76%) | 103 (84%) | 71 (85%) | 41 (71%) | 60 (57%) |
| Male | 209 (24%) | 30 (33%) | 117 (22%) | 74 (18%) | 0 (0%) | 33 (15%) | 36 (24%) | 19 (16%) | 13 (15%) | 17 (29%) | 45 (43%) |
| BMI, kg/m<sup>2</sup> | 25.2 (5.8) | 24.8 (4.3) | 25.1 (6.0) | 25.2 (6.3) | 25.3 (6.5) | 25.8 (6.1) | 24.8 (5.5) | 30.4 (7.3) | 25.5 (5.5) | 25.0 (6.1) | 24.2 (6.3) |
| Race/Ethnicity |  |  |  |  |  |  |  |  |  |  |  |
| White | 636 (73%) | 70 (78%) | 400 (74%) | 298 (74%) | 185 (74%) | 157 (72%) | 112 (75%) | 85 (70%) | 57 (68%) | 44 (76%) | 76 (72%) |
| Hispanic or Latino | 83 (9.5%) | 5 (5.6%) | 59 (11%) | 40 (10.0%) | 29 (12%) | 28 (13%) | 13 (8.7%) | 15 (12%) | 10 (12%) | 3 (5.2%) | 13 (12%) |
| Asian | 59 (6.7%) | 4 (4.4%) | 38 (7.0%) | 29 (7.2%) | 15 (6.0%) | 14 (6.5%) | 11 (7.3%) | 10 (8.2%) | 7 (8.3%) | 6 (10%) | 5 (4.8%) |
| Black or African American | 11<br>(1.3%) | 2<br>(2.2%) | 9<br>(1.7%) | 9<br>(2.2%) | 5<br>(2.0%) | 1<br>(0.5%) | 0<br>(0%) | 2<br>(1.6%) | 2<br>(2.4%) | 0<br>(0%) | 0<br>(0%) |
| Multiple | 37<br>(4.2%) | 3<br>(3.3%) | 26<br>(4.8%) | 18<br>(4.5%) | 10<br>(4.0%) | 11<br>(5.1%) | 11<br>(7.3%) | 8<br>(6.6%) | 3<br>(3.6%) | 2<br>(3.4%) | 8<br>(7.6%) |
| Other | 24<br>(2.7%) | 6<br>(6.7%) | 12<br>(2.2%) | 7<br>(1.7%) | 6<br>(2.4%) | 6<br>(2.8%) | 3<br>(2.0%) | 2<br>(1.6%) | 5<br>(6.0%) | 3<br>(5.2%) | 3<br>(2.9%) |
| Unknown | 27<br>(3.1%) | 0<br>(0%) | 0<br>(0%) | 0<br>(0%) | 0<br>(0%) | 0<br>(0%) | 0<br>(0%) | 0<br>(0%) | 0<br>(0%) | 0<br>(0%) | 0<br>(0%) |
| Occult Blood |  |  |  |  |  |  |  |  |  |  |  |
| Negative | 832<br>(95%) | 83<br>(92%) | 519<br>(95%) | 387<br>(97%) | 245<br>(98%) | 207<br>(95%) | 145<br>(97%) | 114<br>(93%) | 78<br>(93%) | 53<br>(91%) | 99<br>(94%) |
| Positive | 45<br>(5.1%) | 7<br>(7.8%) | 25<br>(4.6%) | 14<br>(3.5%) | 5<br>(2.0%) | 10<br>(4.6%) | 5<br>(3.3%) | 8<br>(6.6%) | 6<br>(7.1%) | 5<br>(8.6%) | 6<br>(5.7%) |
| Fecal Calprotectin |  |  |  |  |  |  |  |  |  |  |  |
| In range | 795<br>(91%) | 78<br>(87%) | 489<br>(90%) | 368<br>(92%) | 234<br>(94%) | 201<br>(93%) | 137<br>(91%) | 108<br>(89%) | 74<br>(88%) | 45<br>(78%) | 88<br>(84%) |
| High (≥80 µg/g) | 81<br>(9.2%) | 12<br>(13%) | 55<br>(10%) | 33<br>(8.2%) | 16<br>(6.4%) | 16<br>(7.4%) | 13<br>(8.7%) | 14<br>(11%) | 10<br>(12%) | 13<br>(22%) | 17<br>(16%) |

Data are presented as mean (standard deviation) for continuous variables (Age, BMI) and as count (percentage) for categorical variables. The "Self-reported conditions" categories are not mutually exclusive, as participants could report conditions from multiple categories. Percentages are shown for the total cohort (N=877) and for each condition category. SD, standard deviation; BMI, body mass index; IBD, inflammatory bowel disease.

Most participants had stool chemistry metrics within the reference range (**Fig. 1c; Suppl. File 1, Table S3**). Elevated calprotectin levels were observed in 9.4% of participants. Occult blood was detected in 5.1% of samples, including some from participants who self-identified as healthy (**Table 1**).

Before examining microbiome associations, we first evaluated relationships among the stool chemistry metrics themselves. As expected, SCFA percentages showed strong compositional relationships, with acetate percentage inversely correlated with butyrate percentage (Spearman *ρ* = −0.69). Stool pH was inversely associated with butyrate (Spearman *ρ* = −0.35). In contrast, inflammatory and immune-related metrics formed a correlated cluster: calprotectin and lactoferrin were positively associated (Spearman *ρ* = 0.31) and were frequently elevated together (Fisher’s exact test odds ratio ≈ 101 for joint out-of-range status, *q* < 0.001), with secretory IgA and lysozyme showing a similar pattern (**Suppl. File 1**, **Tables S4 and S5**; **Suppl. File 2**, **Fig. S1 and S2**).

### Stool chemistry metrics correspond to shifts in microbiome diversity and community structure

To explore how gut microbiome diversity and composition are associated with stool chemistry metrics, we characterized microbiome alpha diversity (overall richness and evenness), beta diversity (Bray-Curtis dissimilarity, differences in overall microbiome community composition between individuals and between groups). We then evaluated the robustness of these associations using models adjusted for potential confounding factors. We first assessed stool chemistry metrics related to inflammation and immunity. Calprotectin (**Fig. 2a**) and lactoferrin (**Suppl. File 2, Fig. S3)** were both associated with shifts in beta diversity (PERMANOVA *p* = 0.001 and *p* = 0.002, respectively; PERMDISP2, *p* = 0.002 for both), but not alpha diversity. However, each explained relatively small fractions of the variance of the microbiome (0.41% and 0.3%, respectively; **Fig. 2b**). Meanwhile, individuals with sIgA levels below the reference range showed significantly higher alpha diversity (*q* < 0.001; **Fig. 2a**). sIgA was also associated with differences in beta diversity (PERMANOVA *p* = 0.001), driven by greater dispersion among individuals with low sIgA levels (PERMDISP2, *p* = 0.007; **Fig. 2a**), and explained 0.5% of the variance in microbiome composition (**Fig. 2b**).

**Fig. 2:**
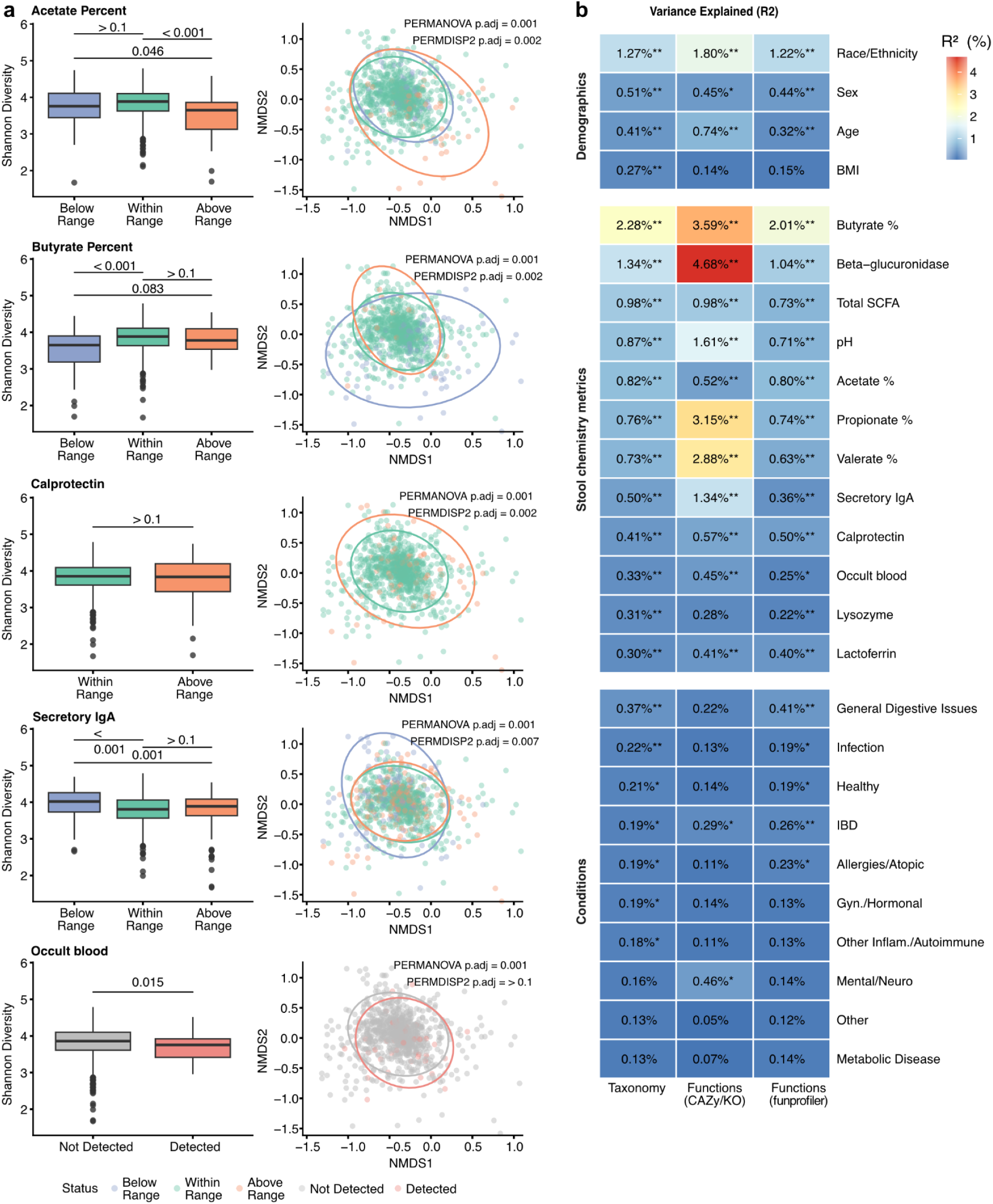
Stool chemistry metrics are associated with gut microbiome diversity, composition, and variance explained across taxonomic and functional profiles. **(a)** Shannon diversity (left panels) and non-metric multidimensional scaling (NMDS; right panels) plots showing gut microbiome variation according to stool chemistry metrics, including acetate percent, butyrate percent, calprotectin, secretory IgA, and occult blood. Samples were grouped as below range (blue), within range (green), or above range (orange) based on reference ranges for each metric; except for occult blood in which samples were grouped as not detected (grey) or detected (red). Boxplots display Shannon diversity indices, with statistical differences assessed using the Wilcoxon rank-sum test and Benjamini–Hochberg–adjusted *p*-values (*q*-values) indicated above group comparisons. NMDS ordinations are based on Bray–Curtis dissimilarities, with ellipses indicating 95% confidence intervals. Statistical significance in community composition was tested using PERMANOVA, and differences in dispersion were evaluated using PERMDISP2. Two outlier values were omitted from these plots to improve visualization; all statistical analyses were performed on the full dataset. **(b)** Heatmap showing the proportion of variance explained (R²) by demographic variables, stool chemistry metrics, and health conditions across three data types: taxonomic composition, targeted functional profiles (KO/CAZy), and inferred functional profiles (FunProfiler). Warmer colors indicate higher explanatory power. Asterisks denote statistical significance (* *p* < 0.05; ** *p* < 0.01).

The presence of occult blood was associated with lower alpha diversity (*q* = 0.015), and beta diversity differed between individuals with and without occult blood in their stool (PERMANOVA *p* = 0.001; PERMDISP2, *p* > 0.1; **Fig. 2a**). Overall, presence of occult blood explained 0.33% of the variance of the microbiome (**Fig. 2b**). Because occult blood was detected in seven self-reported healthy participants (**Table 1**), we investigated this finding further. Occult blood positivity was approximately threefold more common in males than in females (10.5% [22/209] vs. 3.4% [23/668]; **Suppl. File 1, Table S6**). Among the seven occult blood-positive self-reported healthy participants, six were male. Similar patterns of alpha and beta diversity were observed within each sex and, despite the limited sample size, within the self-reported healthy subset (Shannon diversity: 3.81 vs. 4.09, *p* = 0.030; PERMANOVA R² = 0.0176, *p* = 0.026; **Suppl. File 1, Table S6**).

We then examined SCFAs. Individuals with acetate percentages above the reference range showed significantly lower alpha diversity than those within range (**Fig. 2a**; *q* < 0.001). They also showed changes in overall microbiome composition, with greater differences between individuals in the group—that is, higher beta-diversity dispersion, indicating their microbiomes varied more widely from one another (**Fig. 2a**; PERMANOVA, *p* = 0.001, PERMDISP2, *p* = 0.002). Acetate percentage explained 0.82% of the variance in the taxonomy of the microbiome (**Fig. 2b**). Individuals with butyrate percentages below the reference range also had significantly lower alpha diversity than those within range (**Fig. 2a**; *q* < 0.001). Their microbiomes grouped separately from the other individuals in the beta diversity analyses, indicating a different overall community structure, and they also showed greater variability within the groups (**Fig. 2a**; PERMANOVA *p* = 0.001, PERMDISP2, *p* = 0.002). Butyrate percentage explained 2.28% of the variance in the taxonomy of the microbiome (**Fig. 2b**). Similar to acetate, individuals with propionate percentages above the reference range showed significantly lower alpha diversity than those within range (*q* < 0.001; **Suppl. File 2, Fig. S3**). Their microbiomes grouped separately from the other individuals (PERMANOVA *p* = 0.001; PERMDISP2, *p* > 0.1; **Suppl. File 2, Fig. S3**), and propionate percentage explained 0.76% of variation in the taxonomy of the microbiome (**Fig. 2b**). Total SCFA concentration was not associated with changes in alpha diversity. However, the overall microbiome composition still differed across individuals with total SCFA concentration below, above, and within the reference range, as shown by beta diversity analyses (PERMANOVA *p* = 0.001; PERMDISP2, *p* = 0.002; **Suppl. File 2, Fig. S3**). Total SCFA concentration explained 0.98% of the variance in the microbiome (**Fig. 2b**).

Other stool chemistry metrics included stool pH and beta-glucuronidase. Individuals with low (more acidic) stool pH had significantly lower alpha diversity (*p* = 0.012; **Suppl. File 2, Fig. S3**); while those with high (more alkaline) pH had microbiomes that differed clearly from the other groups based on beta diversity separation (PERMANOVA *p* = 0.001; PERMDISP2, *p* > 0.1; **Suppl. File 2, Fig. S3**). Stool pH explained 0.87% of variance in the taxonomy of the microbiome (**Fig. 2b**). Individuals with low beta-glucuronidase activity had significantly lower alpha diversity (*p* < 0.001; **Suppl. File 2, Fig. S3**), and beta-glucuronidase explained 1.34% of the variance in the microbiome (**Fig. 2b**).

To assess whether stool chemistry associations were consistent across independent functional profiling approaches, we evaluated both assembly-based, targeted functional annotations and k-mer–based KEGG Orthology (KO) profiles. Functional profiles generated by the two approaches showed substantial concordance among shared KO features (Spearman *ρ* = 0.771, *p* < 0.001), with 62% of individual KOs showing significant positive correlations after BH-FDR correction (*q* < 0.05) (**Suppl. File 1, Table S7**).

In addition to these stool metrics, the key demographics age, sex, BMI, and race each explained a portion of the variance in the taxonomy of the microbiome, at 0.41%, 0.51%, 0.27%, and 1.3%, respectively (**Fig. 2b**). Health conditions also explained a portion of variance in the microbiome, and will be discussed in a later section.

To determine whether these stool chemistry–microbiome associations were independent of demographic, lifestyle, dietary, clinical, and technical confounders, we repeated the variance partitioning analysis while adjusting for age, sex, BMI, race, smoking and alcohol use, estimated dietary fiber intake, probiotic use, medications, stool consistency and estimated transit time, sequencing batch, and sequencing depth. All 12 stool chemistry metrics remained significantly associated with microbiome taxonomy after adjustment (*q* = 0.005), and 11 of 12 remained associated with targeted functional gene profiles, with lysozyme as the only exception (*q* ≤ 0.08). All 12 also remained significant against the untargeted KO profiles (*q* ≤ 0.01; **Suppl. File 1, Table S8; Suppl. File 2, Fig. S4**). Adjustment resulted in only modest reductions in the proportion of explained variance for most metrics. For example, the variance explained by butyrate percentage decreased from 2.3% to 1.8%, while propionate showed little change. Total SCFA concentration exhibited the greatest attenuation but remained significantly associated with both taxonomy and functional profiles (**Suppl. File 1, Table S8**). Across all evaluated covariates, race accounted for the largest proportion of unique microbiome variation (1.1% for taxonomy and 1.1-1.6% for functions), whereas others like estimated dietary fiber and stool transit time contributed less than 1% (**Suppl. File 1, Table S9**).

### Associations between self-reported conditions, microbiome, and stool chemistry metrics

Next, we wanted to see if self-reported health conditions correlated with changes in microbial diversity, stool chemistry metrics, or taxonomic profiles. Individuals reporting either IBD or general digestive issues displayed significantly lower alpha diversity compared with healthy participants (**Suppl. File 2, Fig. S5**). Beta diversity analyses further revealed clear differences in community structure: healthy individuals formed a tightly clustered group, whereas both the general digestive issues group and the IBD group exhibited distinct and highly variable compositional shifts (**Suppl. File 2, Fig. S5**). Despite these compositional shifts, having general digestive issues or IBD explained only a small fraction of the variation in the taxonomic profiles of the microbiome, with each condition accounting for less than 0.5% of the variance (**Fig. 2b**).

To identify condition-specific microbial signatures, we applied Multivariable Association with Linear Models 2 (MaAsLin2)^12^ to test associations between species-level relative abundances and disease groupings (FDR < 0.05). In the IBD group, enrichment was observed for *Citrobacter*, *Klebsiella*, and *Peduovirus*. Participants reporting general digestive symptoms exhibited increased abundances of *Escherichia coli*, *Escherichia flexneri*, *Clostridium_AP scindens*, and *Peduovirus* (**Suppl. File 1, Table S10**).

Analysis of stool chemistry metrics showed that lactoferrin levels were elevated in IBD relative to individuals with other conditions; however, this association did not remain significant after correction for multiple comparisons (**Suppl. File 1, Table S11**). Notably, several participants who self-reported as healthy tested positive for occult blood, or exhibited elevated calprotectin or lactoferrin levels (**Suppl. File 1, Table S11**), suggesting that self-reported health status may not reliably capture underlying intestinal inflammation or mucosal damage. To assess the impact of this potential misclassification, we compared the self-reported healthy group with a stool chemistry-defined healthy subgroup consisting of participants with within range calprotectin and lactoferrin measurements and no detected occult blood. Of the 90 participants who self-reported as healthy, 21.1% (n = 19) had at least one abnormal metric (**Suppl. File 1, Table S12**). These stool chemistry-discordant participants exhibited lower Shannon diversity than the stool chemistry-defined healthy subgroup (median 3.92 vs. 4.17; *p* = 0.085; rank-biserial correlation = 0.26). Using the stool chemistry-defined healthy subgroup (n = 71) as the reference strengthened all healthy-versus-condition comparisons while applying the same age-, sex-, and BMI-adjusted models. For example, in IBD, the adjusted alpha diversity coefficient increased from 0.200 to 0.255 (*p* = 0.018 to 0.002), and the unique adjusted beta diversity *R*² increased from 0.013 to 0.018. Similar increases in effect size were observed for general digestive issues and metabolic disease (**Suppl. File 1, Table S13**). These findings indicate that inclusion of stool chemistry-abnormal participants within the self-reported healthy group attenuated, rather than exaggerated, the observed microbiome differences.

Additionally, a sensitivity analysis was performed to assess the robustness of associations between stool chemistry categories and microbial diversity metrics after excluding participants with IBD. Alpha and beta diversity effect sizes were largely consistent between the full cohort and the No-IBD cohort, indicating that the observed diversity associations were not driven by IBD (**Fig. 3a, b**). Excluding individuals with IBD resulted in modestly larger effect sizes for select SCFA-related comparisons, particularly butyrate and propionate percentages, while all other metrics were largely unchanged. No associations were substantially attenuated after IBD exclusion. This pattern is consistent with the uniformly reduced Shannon diversity observed in individuals with IBD (**Fig. 3c**), which may attenuate between-group contrasts when included, rather than reflecting differences in SCFA distributions, which were largely within the reference range.

**Fig. 3:**
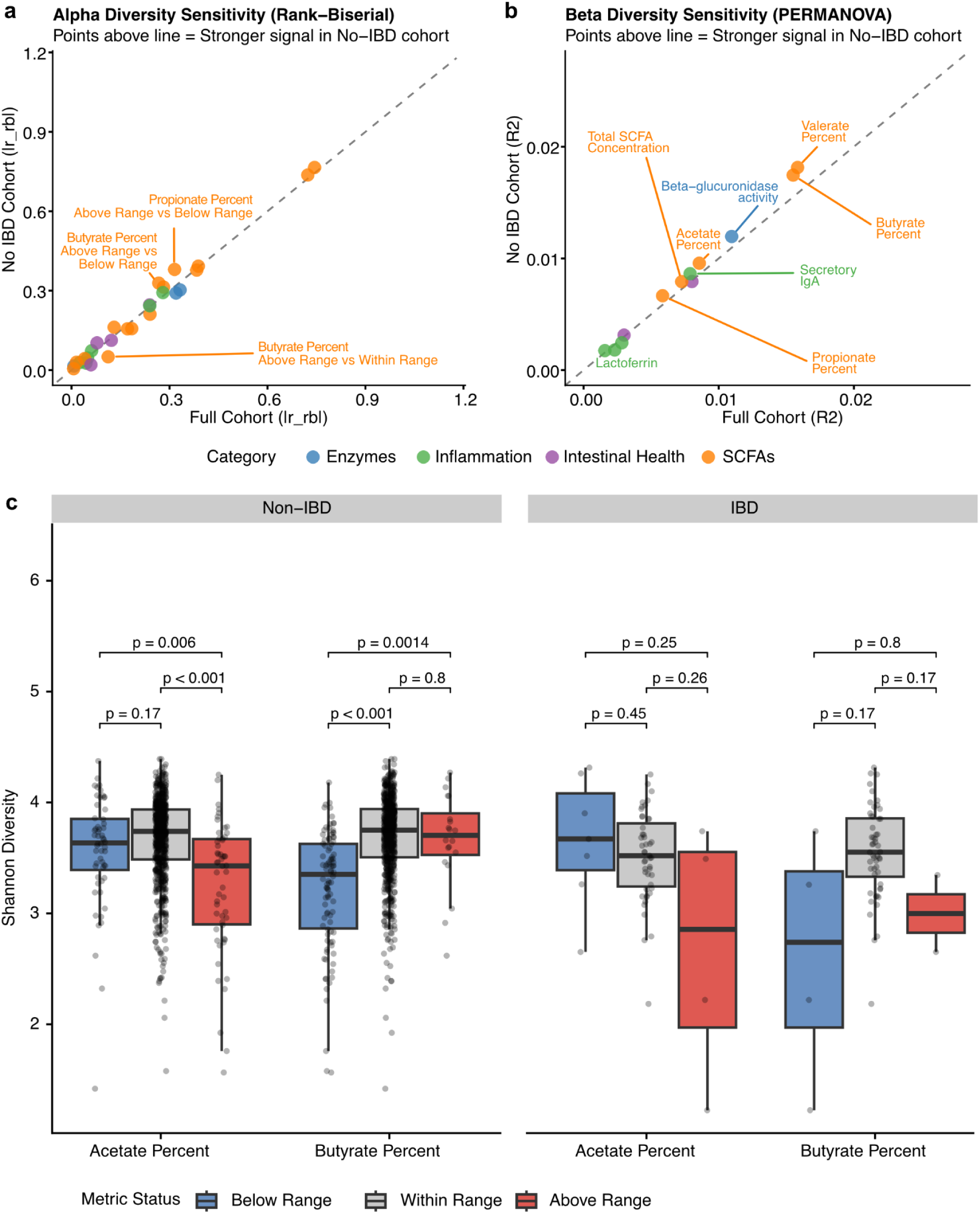
Sensitivity analyses show that stool chemistry–microbiome associations are largely robust to exclusion of inflammatory bowel disease. **(a)** Alpha diversity sensitivity analysis, effect sizes (as measured by Glass’ rank-biserial correlation) are shown for the full cohort (x-axis) versus the No-IBD cohort (y-axis). Each point represents a stool chemistry metric comparison and is colored by functional category. Points along the diagonal indicate comparable effect sizes between analyses, whereas points above the diagonal indicate stronger associations in the No-IBD cohort. **(b)** For beta diversity, PERMANOVA R² values are similarly compared between cohorts. **(c)** Sensitivity analysis of alpha diversity associations for acetate percent and butyrate percent after stratification by IBD status. Boxplots show Shannon diversity for samples with Below Range (blue), Within Range (grey), or Above Range (red) acetate or butyrate percentages, based on established reference ranges. Results are shown separately for participants without IBD (left) and with IBD (right). Pairwise differences between categories were assessed using the Wilcoxon rank-sum test, and Benjamini–Hochberg-adjusted *p*-values are shown above each comparison.

Together, these findings show that disruptions in gut health are accompanied by reduced microbial diversity, altered community structure, and condition-specific taxonomic shifts, while inflammatory stool chemistry abnormalities can be present even in individuals who self-report as healthy. Furthermore, the associations between stool chemistry metrics and microbiome diversity remained robust after excluding participants with IBD and adjusting for various confounding factors, indicating that these relationships reflect broader links between stool chemistry and microbial community structure.

### Stool chemistry profiles reveal distinct species-level microbial signatures

We applied MaAsLin2 to identify microbial taxa linked to inflammation or SCFA production, highlighting significant associations between species-level relative abundances and stool chemistry metrics (*q* < 0.05). Each species was categorized based on its presumed relationship to the host: health-associated, disease-associated, variable (both health- and disease-associated), and not-categorized (**Suppl. File 1, Table S14**).

Inflammatory chemistry metrics displayed few and weak microbial associations (**Fig. 4b**). Calprotectin, lactoferrin, and sIgA showed mainly negative associations with health-associated taxa (e.g., *Blautia* species), and positive associations with disease-associated taxa (e.g., *Escherichia albertii*) (**Suppl. File 1, Tables S14 and S15**). Lysozyme exhibited the fewest significant associations and the lowest cumulative coefficients.

**Fig. 4:**
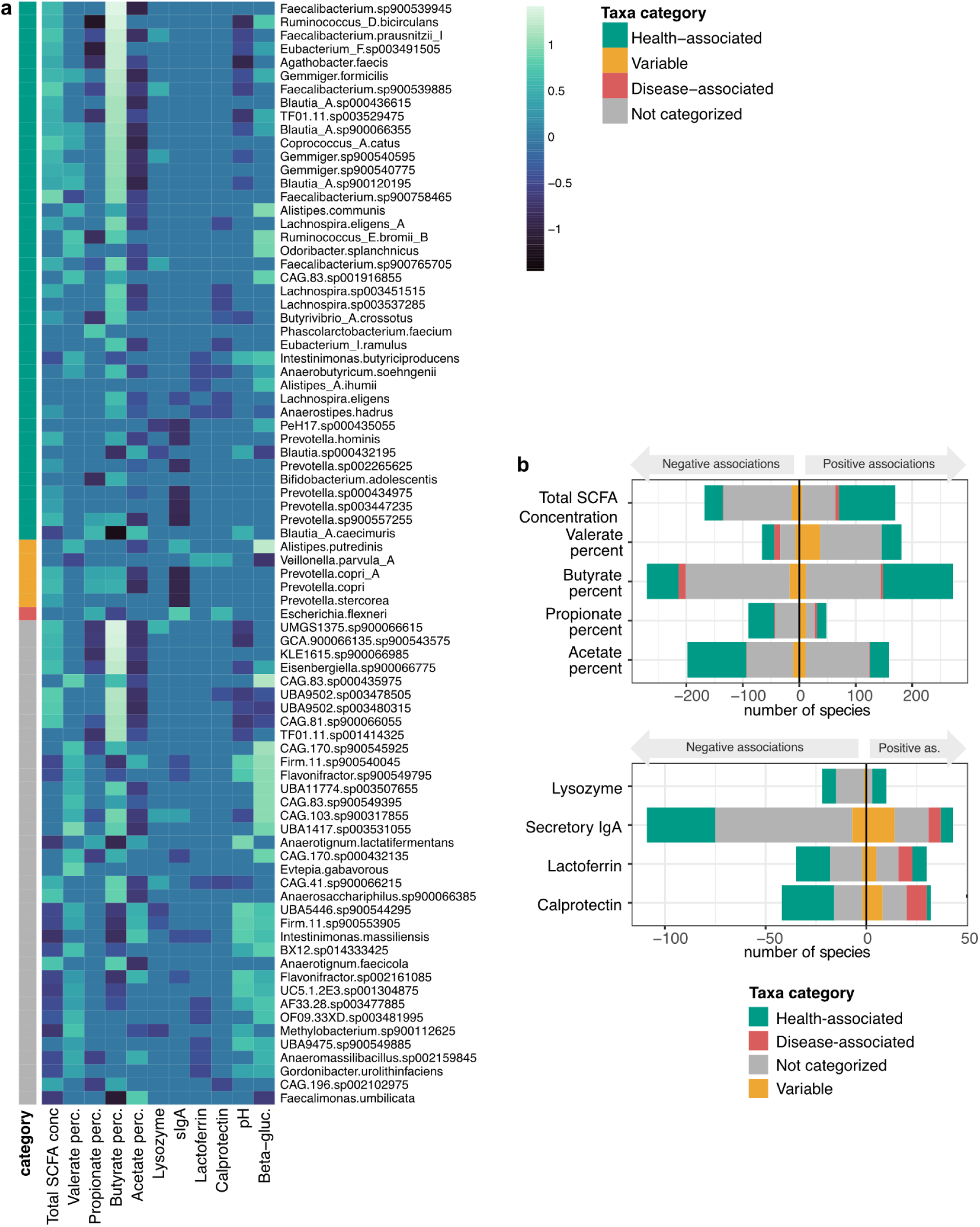
Health-associated and uncategorized taxa dominate SCFA associations, while stool chemistry metrics related to inflammation show fewer microbial links. Microbial species-level associations with stool chemistry metrics were identified using MaAsLin2. Taxa were categorized based on predefined host-health relationships: Health-associated (green), disease-associated (red), variable (orange), or not-categorized (gray) (Suppl. File 1, Table S14). **(a)** Heatmap displaying microbial species (rows) and stool chemistry metrics (columns). To highlight the most robust signatures, only the top 10 species per metric, ranked by absolute MaAsLin2 coefficient, are shown. Species are grouped by health category. The color gradient denotes the direction and strength of the association, with light teal indicating positive and dark purple indicating negative associations. All displayed cells represent statistically significant associations (FDR-corrected *q* < 0.05). (**b**) Total count of significant microbial associations per metric (*q* < 0.05). Bars represent the cumulative number of species within each category showing positive (above zero) or negative (below zero) associations.

For total SCFA concentration, most of the significant positive associations were linked to health-associated taxa, such as *Faecalibacterium* species (**Suppl. File 1, Tables S14 and S15**). In contrast, butyrate percentage displayed a more balanced distribution, with both positive and negative associations with health-associated taxa (e.g., *Faecalibacterium* species and *Blautia_A caecimuris*, respectively). For acetate and propionate, the predominant pattern was negative associations with health-associated taxa such as *Ruminococcus_D bicirculans* (GTDB r.232: *Hominimerdicola aceti*)*, Eubacterium_F sp003491505*, and *Faecalibacterium sp900539945* (**Fig. 4a; Suppl. File 1, Tables S14 and S15**), suggesting that higher acetate and propionate percentages are associated with a lower relative abundance of beneficial butyrate producers. Because SCFA percentages are compositional, an increase in acetate or propionate necessarily corresponds to a lower percentage of butyrate.

Finally, stool pH and β-glucuronidase activity showed heterogeneous associations across all taxa categories (**Suppl. File 1, Table S14**). Neither showed a clear taxonomic pattern, indicating that these metrics may be influenced by a wider range of microbial and host factors (**Suppl. File 1, Tables S14 and S15**).

Alongside shifts in bacteria with well-characterized roles in gut health, some chemistry metrics were also associated with significant shifts in less characterized bacteria. For example, the species *Anaerococcus vaginalis* and *Peptostreptococcus anaerobius* were both associated with increased calprotectin, lactoferrin, and sIgA, and lower butyrate percentage (**Suppl. File 1, Table S14**). Species such as *Anaerobutyricum sp900554965* (GTDB r.232: *Anaerobutyricum hallii*)*, CAG-41 sp900066215* (GTDB r.232: *Hominilimicola fabiformis*), and *Phocaeicola sp900553715* (GTDB r.232: *Phocaeicola coprocola*) were each associated with increased butyrate, and decreased calprotectin and lactoferrin (**Suppl. File 1, Table S14**), a pattern consistent with health-associated bacteria. Similarly, the recently isolated species *Intestinimonas massiliensis* was associated with increased acetate, and decreased lactoferrin and sIgA (**Suppl. File 1, Table S14**).

To evaluate the robustness of these associations, we progressively adjusted MaAsLin2 models for demographic, lifestyle, dietary, clinical, and technical covariates. Across all stool chemistry metrics and taxonomic levels, no association changed direction after adjustment (**Suppl. File 1, Table S16**). Although adjustment reduced the number of significant associations, nearly half or more of the SCFA-taxon associations remained significant in the fully adjusted models (species/genus: butyrate percentage 64%/76%, propionate 54%/64%, acetate 49%/61%; **Suppl. File 1, Table S17**). Demographic covariates accounted for the largest reduction in significant associations, whereas adjustment for estimated dietary fiber intake and stool transit had relatively little impact, indicating that these factors did not explain the observed SCFA-microbiome relationships. In contrast to the SCFAs, inflammation-related metrics showed an initial increase in significant associations after demographic adjustment before declining with further adjustment. For example, calprotectin and lysozyme exhibited more significant associations after demographic adjustment than in the minimally adjusted model, suggesting that demographic factors had initially masked some microbiome associations (**Suppl. File 1, Table S18**).

The robustness of stool chemistry associations extended to functional gene profiles. Applying the same progressive adjustment framework to both targeted assembly (PLASS; 264 genes) and untargeted functional profiling (funprofiler; 9,144 KOs) yielded patterns consistent with the taxonomic analyses. Across both pipelines, no functional gene association changed direction after adjustment, and 37–87% of significant associations remained in the fully adjusted models, with higher retention observed for PLASS than funprofiler (**Suppl. File 1, Table S19**). As in the taxonomic analyses, demographic covariates accounted for the largest reduction in significant associations, whereas estimated dietary fiber intake and stool transit had comparatively little impact (**Suppl. File 1, Table S20**).

Given the female predominance of the cohort (76%), we performed a sex-stratified sensitivity analysis to evaluate the robustness of stool chemistry-microbial associations. Fully adjusted MaAsLin2 models were fit separately in females (n = 668) and males (n = 209). Associations that reached significance in both sexes were directionally consistent across the subset of stool chemistry metrics evaluated (**Suppl. File 1, Table S21**). Similarly, adjusted PERMANOVA effect sizes were comparable between females and males, with similar variance explained for butyrate percentage (R² = 0.0182 and 0.0176, respectively), while pH and sIgA explained comparable or greater variance in males. Although fewer significant associations were identified in males, this likely reflects reduced statistical power due to the smaller male sample size rather than sex-specific differences in the underlying associations (**Suppl. File 1, Table S22**).

Together, these findings show that total SCFA concentration is predominantly associated with health-associated taxa, whereas inflammatory stool chemistry metrics show weaker associations primarily with disease-associated species. Several not-categorized taxa exhibited association patterns that suggest either pro-inflammatory or health-associated roles, warranting further investigation into their functional contributions to gut health. The observed associations remained directionally consistent after extensive confounder adjustment, were also observed in functional gene analyses, and were comparable across sexes, supporting the robustness of the observed microbiome–stool chemistry relationships.

### Microbiome networks are associated with key stool chemistry patterns

To visualize relationships between gut microbial features and stool chemistry, we constructed a bipartite network of significant associations between microbial taxa and stool chemistry metrics. This revealed clusters of taxa that co-varied with specific metrics, providing insight into microbiome patterns linked to gut health and dysbiosis. To simplify interpretation, we centered the network on butyrate, a major microbial metabolite involved in colonocyte metabolism and immune regulation, together with calprotectin and lactoferrin, two markers of intestinal immune activation. Two distinct neighborhoods emerged: one associated with a healthy, non-inflammatory gut state (increased butyrate, decreased calprotectin and lactoferrin) and another consistent with dysbiosis and inflammation (decreased butyrate, increased calprotectin and lactoferrin) (**Fig. 5**).

**Fig. 5:**
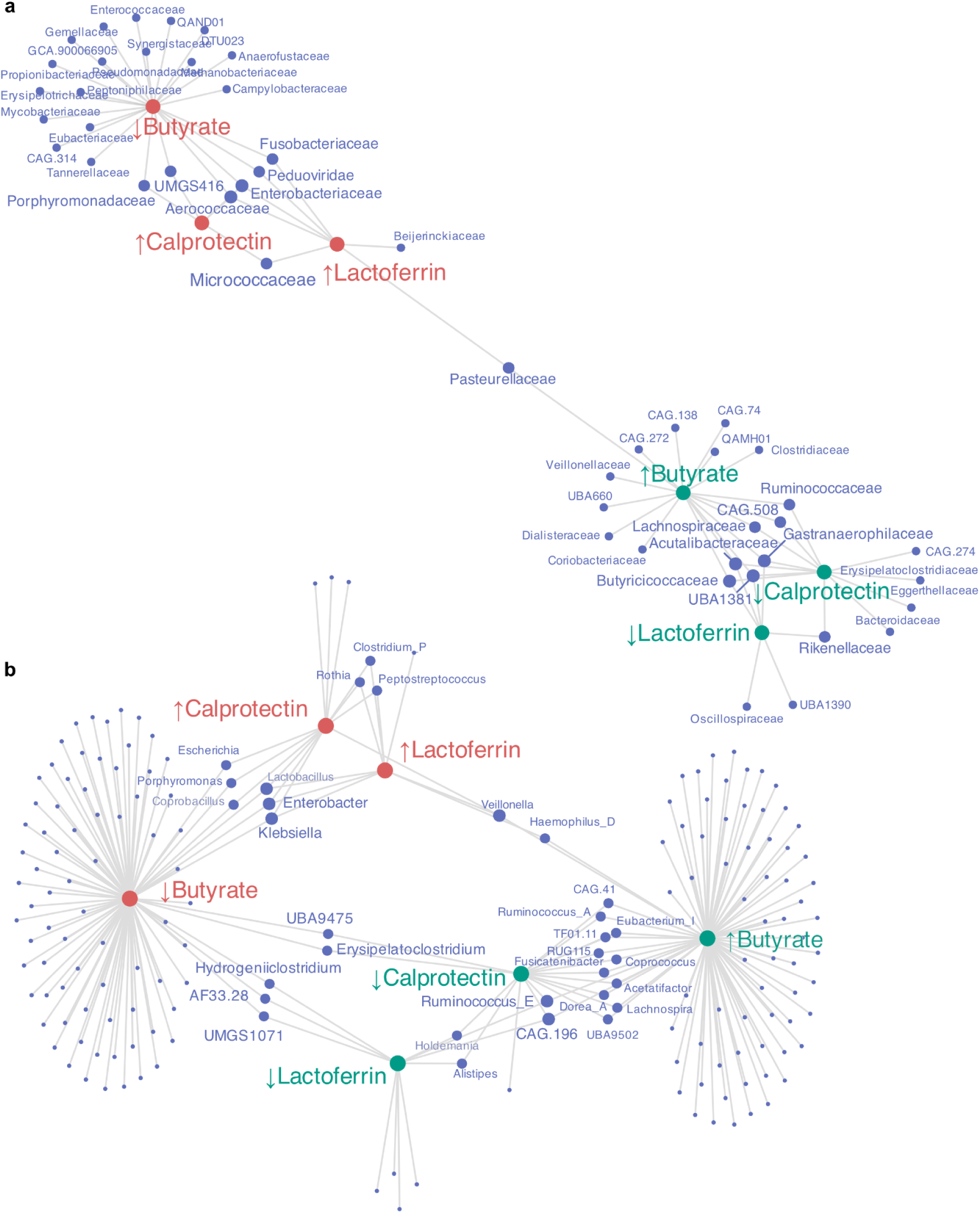
Network of significant associations between select stool chemistry metrics and microbial families and genera. Nodes represent microbial families **(a)** or genera **(b)** and stool chemistry metrics, which were split into “increased” and “decreased” categories. Edges indicate significant associations from MaAsLin2 (FDR q < 0.05). Node size corresponds to the number of significant associations (degree), with larger nodes highlighting features connected to multiple relationships. Colors distinguish microbial features (blue) from stool biomarkers (red and green). The network is limited to features associated with butyrate, calprotectin, and lactoferrin.

The health-associated neighborhood was dominated by anaerobic Firmicutes, including *Ruminococcaceae* and *Lachnospiraceae*, along with less characterized families such as *Acutalibacteraceae*, *Gastranaerophilaceae*, *Butyricicoccaceae*, UBA1381, and CAG-508 (**Fig. 5a; Suppl. File 1, Table S23**). Within these, genera including *Ruminococcus_E* (GTDB r.232: *Ruminococcoides*), CAG-196, *Acetatifactor*, *Coprococcus*, *Lachnospira*, *Ruminococcus_A (*GTDB r.232: *Oliverpabstia)*, *Eubacterium_I*, *Dorea_A*, *Fusicatenibacter*, and TF01-11 (GTDB r.232: *Butyribacter*) were linked to increased butyrate and decreased inflammatory metrics (**Fig. 5b; Suppl. File 1, Table S23**).

The dysbiosis-associated neighborhood comprised families and genera connected to inflammatory conditions. *Aerococcaceae* and *Enterobacteriaceae* were associated with decreased butyrate and increased calprotectin and lactoferrin, while UMGS416, *Porphyromonadaceae*, *Fusobacteriaceae*, and *Peduoviridae* were linked to decreased butyrate and either increased calprotectin or increased lactoferrin (**Fig. 5a; Suppl. File 1, Table S24**). Genera such as *Klebsiella*, *Enterobacter*, *Escherichia*, *Porphyromonas*, and *Coprobacillus* were key nodes in this cluster (**Fig. 5b; Suppl. File 1, Table S24**). Notably, *Lactobacillus* also appeared here.

Together, these findings demonstrate that gut microbiomes cluster into two distinct network neighborhoods—one dominated by anaerobic, butyrate-producing Firmicutes associated with reduced inflammation, and another characterized by Proteobacteria and facultative anaerobes linked to elevated inflammatory metrics and depleted butyrate—revealing coordinated shifts in microbial community structure that distinguish healthy gut ecosystems from dysbiotic states.

Functional analysis was consistent with these taxonomic patterns. The health-associated neighborhood was enriched for genes involved in complex carbohydrate degradation and spore formation (**Suppl. File 1, Table S23**). In contrast, the dysbiosis-associated neighborhood was enriched for functions related to bacterial respiration (nitrate/nitrite respiration, oxidative phosphorylation, ubiquinone synthesis), virulence (siderophore production), and utilization of host-derived compounds (gluconate) (**Suppl. File 1, Table S24**).

These analyses indicate that health-associated gut microbial communities are dominated by commensal organisms that are anaerobic, fiber-degrading, and spore-forming, whereas dysbiosis-associated communities are enriched for pathobionts–facultative anaerobes or aerobes capable of respiratory metabolism, inflammation-related functions, and potential pathogenicity.

### Microbiome classifiers predict stool chemistry metrics with modest accuracy

To evaluate whether gut microbiome profiles can predict stool chemistry metrics, we trained elastic-net classifiers on taxonomic features for 12 stool chemistry metrics, framing each target as a binary, out-of-range classification. Full results for all evaluable tasks are available in **Suppl. File 1, Table S25**. Our primary analyses focused on four targets with sufficient sample sizes across classification groups (at least 25 samples per group): presence of occult blood, high calprotectin (above reference range), low butyrate percentage (below reference range), and high acetate percentage (above reference range) (**Fig. 6 and Table 2**).

**Fig. 6:**
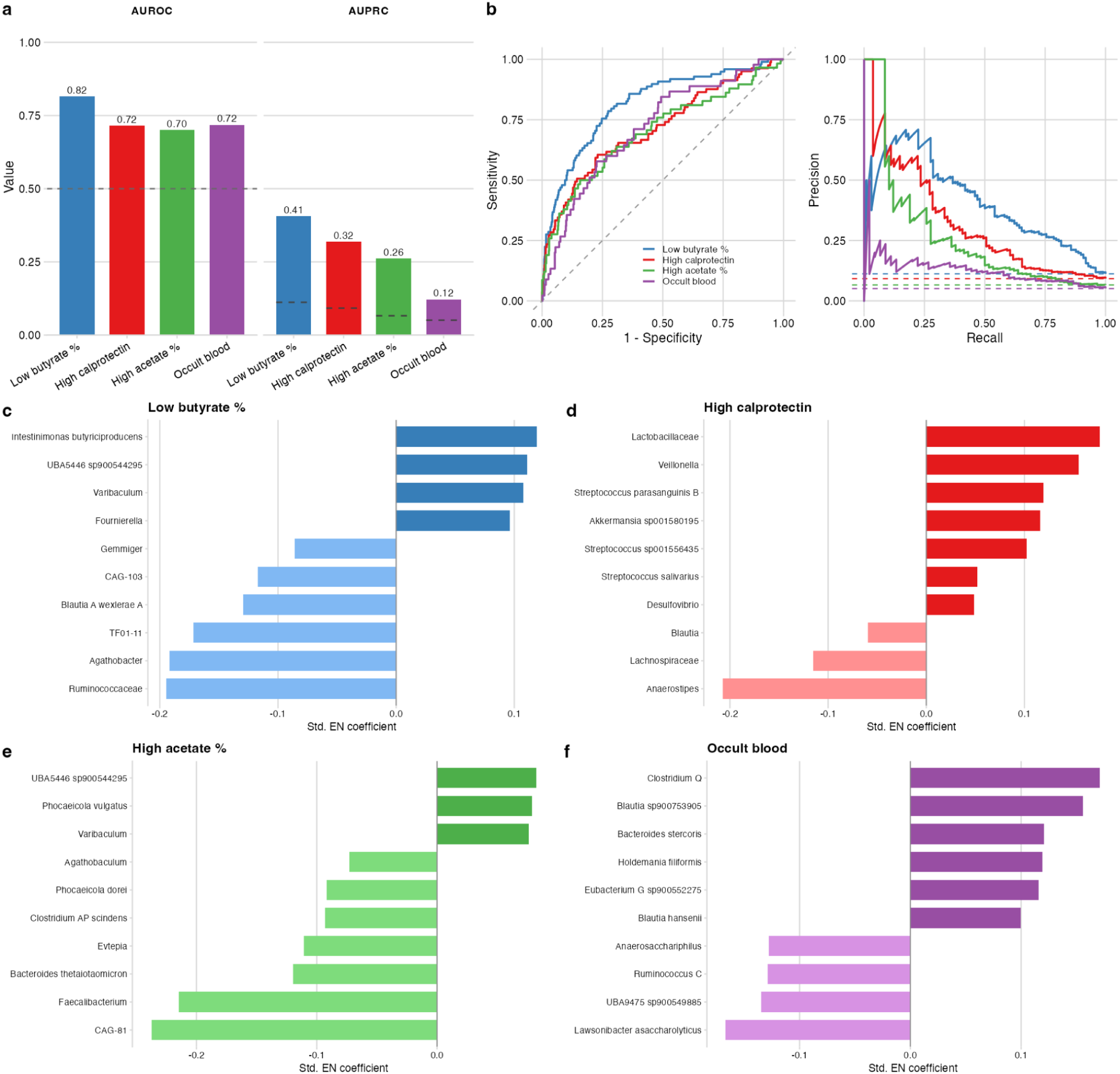
Microbiome-based classifiers identify out-of-range stool chemistry with modest, target-dependent discrimination. Classifiers were elastic-net logistic regressions trained on centered log-ratio–transformed taxonomic features at species, genus and family level (n = 877). Performance was estimated by five-fold cross-validation with folds grouped by account, so that samples from the same household never spanned a train/test split, and stratified by class; feature filtering was applied within training folds only. (a) Discrimination for the four headline targets — low butyrate percentage, high calprotectin, high acetate percentage and occult blood — shown as area under the receiver operating characteristic curve (AUROC) and area under the precision–recall curve (AUPRC). Dashed lines indicate chance performance: 0.5 for AUROC, and the target’s own prevalence for AUPRC (11.2%, 9.2%, 6.6% and 5.1% respectively), since AUPRC baselines are prevalence-dependent and not comparable across targets. (b) Receiver operating characteristic (left) and precision–recall (right) curves for the same four targets, computed from held-out cross-validation predictions. The dashed diagonal and the dashed horizontal lines indicate chance performance. (c–f) The ten highest-magnitude standardized elastic-net coefficients for (c) low butyrate percentage, (d) high calprotectin, (e) high acetate percentage and (f) occult blood. Coefficients are standardized (β × SD of the feature) so that CLR-transformed features are comparable, and are taken from a single elastic-net fit to all 877 samples; these panels are for interpretation of feature direction and are not a performance estimate. Positive coefficients (dark bars) indicate taxa associated with the out-of-range state; negative coefficients (light bars) indicate taxa associated with the in-range state. Full-panel performance for all 19 evaluable classification tasks is given in **Suppl. File 1, Table S25**.

**Table 2.**
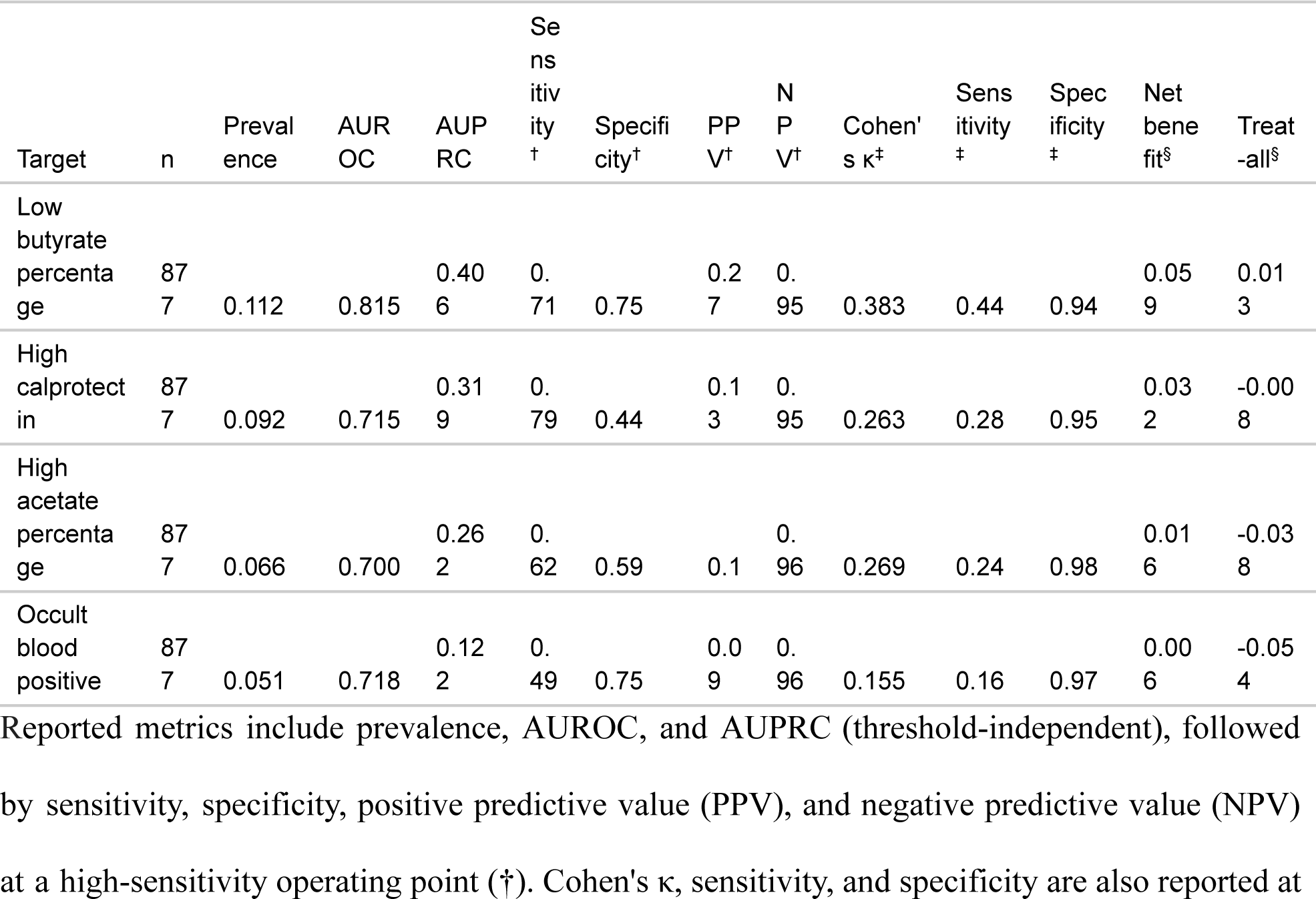
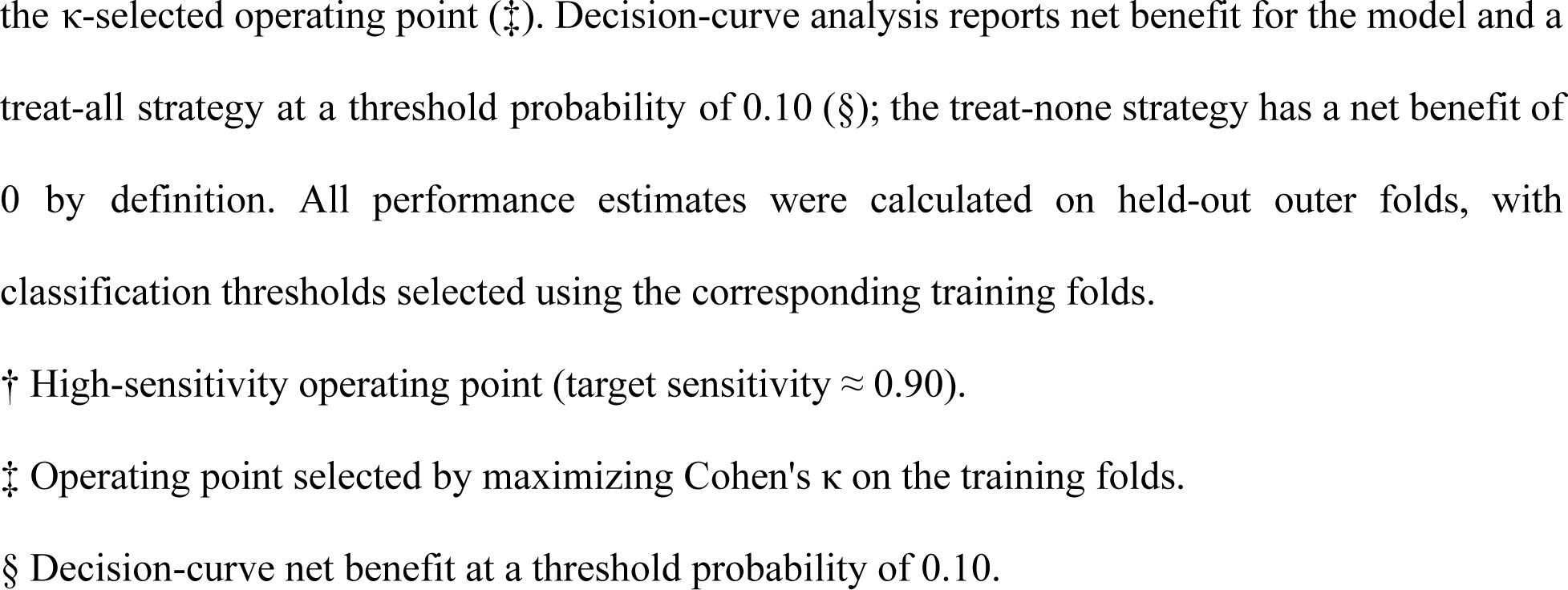
Performance of the four targets with sufficient sample sizes across classification groups.

| Target | n | Prevalence | AUR OC | AUP RC | Sensitivity<sup>†</sup> | Specificity<sup>†</sup> | PPV<sup>†</sup> | NPV<sup>†</sup> | Cohen's $\kappa$<sup>‡</sup> | Sensitivity<sup>‡</sup> | Specificity<sup>‡</sup> | Net benefit<sup>§</sup> | Treat-all<sup>§</sup> |
| --- | --- | --- | --- | --- | --- | --- | --- | --- | --- | --- | --- | --- | --- |
| Low butyrate percentage | 877 | 0.112 | 0.815 | 0.406 | 0.71 | 0.75 | 0.27 | 0.95 | 0.383 | 0.44 | 0.94 | 0.059 | 0.013 |
| High calprotectin | 877 | 0.092 | 0.715 | 0.319 | 0.79 | 0.44 | 0.13 | 0.95 | 0.263 | 0.28 | 0.95 | 0.032 | -0.008 |
| High acetate percentage | 877 | 0.066 | 0.700 | 0.262 | 0.62 | 0.59 | 0.1 | 0.96 | 0.269 | 0.24 | 0.98 | 0.016 | -0.038 |
| Occult blood positive | 877 | 0.051 | 0.718 | 0.122 | 0.49 | 0.75 | 0.09 | 0.96 | 0.155 | 0.16 | 0.97 | 0.006 | -0.054 |

Reported metrics include prevalence, AUROC, and AUPRC (threshold-independent), followed by sensitivity, specificity, positive predictive value (PPV), and negative predictive value (NPV) at a high-sensitivity operating point (<sup>†</sup>). Cohen's $\kappa$ , sensitivity, and specificity are also reported at
the $\kappa$ -selected operating point (‡). Decision-curve analysis reports net benefit for the model and a treat-all strategy at a threshold probability of 0.10 (§); the treat-none strategy has a net benefit of 0 by definition. All performance estimates were calculated on held-out outer folds, with classification thresholds selected using the corresponding training folds.
† High-sensitivity operating point (target sensitivity $\approx 0.90$ ).
‡ Operating point selected by maximizing Cohen's $\kappa$ on the training folds.
§ Decision-curve net benefit at a threshold probability of 0.10.

Performance of the predictive models was assessed using a variety of metrics (**Table 2**). We evaluated discrimination using the area under the receiver operating characteristic curve (AUROC) and the area under the precision-recall curve (AUPRC), the latter of which is more informative for predicting imbalanced classes. Of the four targets, low butyrate percentage was predicted most accurately, achieving an AUROC of 0.815 and an AUPRC of 0.406 (**Fig. 6a,b**). When the decision threshold was adjusted to prioritize high sensitivity (90%), the model retained a specificity of 0.75, indicating that it correctly identified most low-butyrate samples while retaining a relatively low false-positive rate (**Table 2**). Additionally, the low butyrate model provided a net benefit—a measure of decision performance that balances true positives against false positives—of 0.059, compared with 0.013 when all samples were classified as low. Positive predictive value (PPV) was 0.27 and negative predictive value (NPV) was 0.95 (**Table 2**). *Ruminococcaceae* showed a negative association with the low-butyrate state, consistent with this family containing several butyrate-producing taxa (**Fig. 6c**).

Three additional models were evaluated, but did not perform as well as low butyrate. Elevated calprotectin showed weaker discrimination (AUROC 0.715, AUPRC 0.320; **Fig. 6a, b**). At the high-sensitivity threshold, the model achieved a specificity of 0.44, a net benefit of 0.032, and a PPV and NPV of 0.13 and 0.95, respectively (**Table 2**). Elevated calprotectin was positively associated with oral-associated taxa, including *Streptococcus* and *Veillonella*, and negatively associated with SCFA-producing taxa such as *Blautia*, *Lachnospiraceae*, and *Anaerostipes* **(Fig. 6d).** High acetate percentage showed similar discrimination (AUROC 0.701, AUPRC 0.262; **Fig. 6a, b**), with a specificity of 0.59 at the high-sensitivity threshold. The model provided only a modest net benefit (0.016), with PPV and NPV of 0.1 and 0.96, respectively (**Table 2**). *Phocaeicola vulgatus*, a well-established acetate-producer, was among the strongest positive associations with the high-acetate state (**Fig. 6e**). Occult blood showed the weakest overall performance (AUROC 0.718, AUPRC 0.122; **Fig. 6a, b**). At the high-sensitivity threshold, specificity dropped to 0.75 with little net benefit (0.006), with a PPV of 0.09 and an NPV of 0.96 (**Table 2**). *Clostridium_Q* showed the strongest positive association with occult blood positivity (**Fig. 6f**).

Beyond the binary classification models described above, we assessed whether the microbiome could also predict stool chemistry concentrations directly, using regression, across the stool chemistry metrics. The same pattern held: microbial outputs (butyrate percentage R² = 0.34; β-glucuronidase R² = 0.35; valerate percentage R² = 0.35) were predicted with moderate accuracy, while host-derived markers (calprotectin R² = 0.08; lysozyme R² = 0.06; lactoferrin R² = 0.04) were predicted poorly (**Suppl. File 1, Table S26**).

To assess generalizability, we applied the elastic-net without refitting to an independent, external cohort of 500 kits collected after the internal discovery cohort was finalized (**Suppl. File 1, Table S27**; **Suppl. File 2, Fig. S6**). Performance held up well overall for models tested (minimum 10 samples per group), with little loss in discrimination (mean AUROC 0.706 external vs. 0.702 internal). Low butyrate percentage was the strongest external performer (AUROC 0.804, AUPRC 0.398), while occult blood again performed similar to random guessing (AUROC 0.672, AUPRC 0.16). High acetate percentage showed similar external discrimination (AUROC 0.788, AUPRC 0.283), though this was based on few positive cases (n=38). However, its decision threshold did not transfer, as all validation samples were classified positive, making that operating point uninformative (**Suppl. File 1, Table S27**). For the remaining eight targets, sensitivity was preserved at the high-sensitivity operating point (external mean 0.88), but specificity was not (mean 0.34; PPV 0.06–0.38, NPV 0.89–0.97; **Suppl. File 1, Table S27**).

To confirm the taxa-only elastic-net was an appropriate model choice, we compared it against random forest and gradient-boosted trees (LightGBM), compared taxonomy alone against taxonomy combined with the functional views in a single early-fusion matrix, and compared early fusion against a late-fusion stacked ensemble (LightGBM base learners combined by an elastic-net meta-learner). Across the full panel of targets, the elastic-net and LightGBM were effectively tied (elastic-net won 10 tasks, LightGBM 9; mean ΔAUPRC ≈ 0.001), and the taxonomy-only elastic-net matched or exceeded the late-fusion ensemble on the primary targets, including low butyrate percentage (AUPRC 0.41 vs. 0.36) and high calprotectin (0.32 vs. 0.24; **Suppl. File 1, Table S28**). Adding the functional data did not improve elastic-net discrimination on average. Two alternative methods of classification were tested and did not improve on binary tail classification. A three-class below/within/above model was uninformative (Cohen’s κ ≈ 0), and an ordinal cumulative-logit model was uniformly worse than the paired binary classifiers (binary classifiers outperformed in 13 of 14 comparisons; mean ΔAUPRC ≈ −0.065; Kendall’s τ ≈ 0). Below- and above-range microbiome signatures therefore appear to represent distinct states rather than opposite ends of a single continuum, causing the ordinal model to collapse toward the within-range majority (**Suppl. File 1, Table S29**).

Overall, these results suggest that stool metagenomic profiles yield modest predictive power of stool chemistry measurements, with performance dependent on metric: microbially-derived metabolites were more predictable than host-derived inflammatory markers.

## Discussion

This large-scale metagenomic study of 877 adults with self-reported health conditions integrates stool chemistry metrics with microbial community profiles to elucidate how microbial metabolism and host responses interrelate in gut health. We found that deviations in chemistry metrics associated with microbial activity (e.g., SCFAs) and host inflammation (e.g., calprotectin and lactoferrin) corresponded with distinct microbial ecological patterns.

Multiple analyses identified stool butyrate percentage as a key stool chemistry metric associated with microbial ecology. Low butyrate was linked to reduced alpha diversity and higher compositional variability, suggesting an unstable community state lacking key butyrate producers^16^. In contrast, the health-associated neighborhood was dominated by anaerobic, fiber-degrading Firmicutes^17,18^ enriched for carbohydrate degradation genes^19^ and spore formation pathways. Together, these features are consistent with a functionally specialized and resilient microbiome. Spore formation, common in Firmicutes, may confer ecological stability, allowing persistence and recolonization after stress^20^, thus supporting intestinal homeostasis through sustained butyrate production. At the taxonomic level, health-associated bacteria were positively associated with total SCFAs and butyrate, consistent with the beneficial role of butyrate in fueling colonocytes, maintaining anaerobiosis, and modulating immunity^17,21^. Conversely, elevated acetate percentages likely reflect disruption of microbial cross-feeding, as acetate is broadly produced but normally consumed by specialized butyrate producers^22^. Because stool SCFAs represent microbial production minus host absorption, the inverse link between butyrate percentage and inflammatory markers in the network analysis suggests that higher stool butyrate percentage reflects a low-inflammatory state with preserved SCFA absorption.

Calprotectin and lactoferrin were associated with significant shifts in beta diversity but not alpha diversity, suggesting that gut inflammation restructures rather than depletes the microbiome. Inflammation alters the intestinal environment through oxidative stress, antimicrobial peptides, and altered nutrient availability^23^, favoring facultative anaerobes over obligate anaerobes and increasing compositional variability^24^. Accordingly, the dysbiosis-associated neighborhood was enriched in facultative anaerobes, pathobionts, virulence-associated functions, and nitrate respiration pathways, suggesting microbial adaptation to the inflamed gut^24–26^. This model in which gut inflammation and altered microbial metabolism are associated with more heterogeneous, dysbiosis-associated ecological states is supported by the greater compositional variability observed in participants with low butyrate.

Beyond microbial metabolism and inflammation, our analyses identified additional microbial features associated with host physiology. Low sIgA was associated with higher alpha diversity. As sIgA is critical for immune homeostasis by coating and containing microbes^27^, this association suggests reduced containment of specific taxa under conditions of low sIgA, though the mechanistic basis of this relationship remains unclear. We also identified numerous uncategorized taxa associated with stool chemistry metrics, indicating that less characterized members of the gut microbiome may contribute meaningfully to gut health. These findings underscore the need for continued functional and taxonomic characterization of the gut microbiome to better define health-associated microbial networks.

In this work’s predictive modeling framework, the gut microbiome was better suited for identifying deficiencies in microbial metabolic output than for detecting host inflammation, with higher negative predictive values than positive predictive values. Consequently, metagenomic predictive models may function better as rule-out screening tools that complement, rather than replace, traditional stool chemistry testing. Predictive performance may improve with larger datasets containing more abnormal events.

This study has several limitations. First, health conditions, diet, and medication/probiotic use were self-reported without clinical validation, introducing potential reporting bias. Second, this retrospective, provider-referred convenience cohort, with a predominance of females, a wide age distribution, and no socioeconomic data, may limit generalizability. Third, relatively few participants had elevated inflammatory markers, restricting our analysis of highly inflamed states. Additionally, reference ranges for stool chemistry metrics are not universally standardized^28,29^; however, consistent results across continuous outcome models suggest our findings were not driven by specific cutoffs. Finally, while many associations were statistically significant, overall effect sizes were small, suggesting limited clinical value for any single feature. Nonetheless, when considered collectively, these microbiome patterns may offer greater clinical insight than individual taxa alone, though these association analyses cannot assign direct function to individual taxa given potential indirect network interactions.

In conclusion, integrating stool chemistry with shotgun metagenomic profiling provides a more comprehensive view of the relationships between the gut microbiome and host physiology than either approach alone. Although the cross-sectional, exploratory design precludes causal inference, our findings identify robust microbiome-stool chemistry associations that warrant further investigation. Future longitudinal and prospective studies together with complementary causal inference approaches such as Mendelian randomization^11,30^, will be needed to establish temporal relationships and validate the potential clinical utility of the microbiome-based classifiers.

## Methods

### Study cohort and measures

This study utilized de-identified data from a convenience sampling-based adult cohort derived from provider-ordered clinical microbiome testing, with all participants authorizing their data to be used for research by agreeing to the Terms of Service. Participant data included demographics (age, sex, BMI, and race/ethnicity), self-reported health conditions, medication use, probiotic use, alcohol use, household smoking exposure, estimated dietary fiber intake (assessed as the average number of servings of plant-based foods consumed per week), stool consistency (assessed using the Bristol Stool Form Scale), bowel movement frequency (bowel movements per day), whole metagenomic shotgun (WMS) sequencing data (taxonomic and functional profiles), and quantified stool chemistry metrics.

Samples from individuals younger than 18 years or 65 years and older, pregnant, or taking antibiotics at the time of collection were excluded.

### Grouping of self-reported conditions

Participants’ self-reported health conditions were manually reviewed and categorized into distinct groups. To ensure consistency and reproducibility, a set of regular expressions was developed and applied to programmatically sort and count these conditions. For instance, any self-reported mentions of “Crohn’s disease,” “ulcerative colitis,” or “inflammatory bowel disease” were grouped under the broader Inflammatory Bowel Disease (IBD) category. Similarly, conditions such as “gluten intolerance,” “irritable bowel syndrome,” and “frequent constipation” were all assigned to the General digestive issues category. The complete list of self-reported conditions and their corresponding categories can be found in **Suppl. File 1, Table S1**.

### Sample collection

The microbiome stool sample kit included a DNA-free, sterile collection system with an active drying tube (FLOQSwab-ADT; Copan Diagnostics, USA)^31^, written instructions, a biosafety bag, and a prepaid return shipping envelope. Participants were instructed to collect a stool sample by swabbing a soiled piece of toilet paper.

The stool chemistry kit included a collection tray, a spoon, a vial, a biosafety bag, a gel pack, and a prepaid return shipping envelope. Participants were instructed to deposit stool into the tray, use the spoon to collect material from three different areas of the sample, transfer it into the vial, and then freeze the vial overnight along with the gel pack before shipping.

### Shotgun metagenomic sample processing and sequencing

Samples were received by and processed in a CLIA/CAP-certified lab, compliant with the Clinical Laboratory Improvement Amendments (CLIA) and accredited by the College of American Pathologists (CAP). Total DNA was extracted using the DNeasy PowerSoil Pro kit (Qiagen, Germany). After quality control (QC), libraries were prepared using the Nextera kit. Library fragment sizes were then assessed using Agilent TapeStation or Bioanalyzer and quantified by qPCR. All samples and libraries passed QC and were sequenced to at least 10 million reads on a NextSeq Illumina Platform.

Fastp^32^ was used for raw read QC. Paired-end reads were trimmed to remove adaptor sequences and low quality reads (minimum length 100 base pairs, mean quality score 20, maximum number of ambiguous bases [N] 5). Duplicate sequences, specifically optical duplicates, were detected and removed. Host sequences that mapped to human genome assembly GRCh38 were removed, and this human sequence data was not saved. We converted the resulting mapping file to a compressed format and extracted the unmapped reads (non-human host containing). A kmer-based method^33^ was used to provide taxonomic classification (confidence 0.1) for the remaining high-quality reads using a database that contains 111,000+ bacterial and archaeal genomes from the GTDB r202 (April 27, 2021: https://data.gtdb.ecogenomic.org/releases/release202/202.0/) and viral (11,618), fungal (390), and protistan (94) genomes available through NCBI GenBank (accessed August 17, 2021). We estimated relative abundance values for detected taxa. All taxa metrics were calculated using an in-house script based on the species relative abundance.

Protein sequences were *de novo* assembled from the microbial reads with PLASS^34^ and functionally annotated based on a combination of BLAST^35^ and HMMER^36^ searches against a curated database of 302 KEGG Orthology HMMs from GenomeNet and 82 CAZy HMMs and FASTA sequences from dbCAN HMMdb v10 and CAZyDB.09242021.fa, respectively. Metagenomic reads were mapped to the functionally annotated protein sequences using MMseqs2^37^ and reads per kilobase per million mapped reads (RPKM) were calculated using an in-house script. To supplement this targeted functional data pipeline, we also implemented funprofiler^38^, a sourmash^39^ kmer sketch-based tool that estimates the relative abundance of all KEGG KOs from raw sequencing reads.

### Stool chemistry processing and metric quantification

Lysozyme, sIgA, and lactoferrin were all measured via ELISA (lysozyme and sIgA: ImmunDiagnostik; lactoferrin: Techlab). Calprotectin was measured via a diagnostic kit (Bühlmann) on a Beckman Coulter DxC 700AU analyzer (Beckman Coulter). SCFAs were measured by gas chromatography. Occult blood was measured using ColoScreen guaiac qualitative detection (Helena Laboratories). Stool pH was measured using MicroEssential Hydrion pH test paper (Micro Essential Laboratory). Beta-glucuronidase was measured with a Laboratory Developed Test based on a previously published method.

Stool chemistry metrics were categorized as “Below,” “Within Range,” or “Above” based on lab-provided healthy reference ranges; except for occult blood which was categorized as “Not detected” or “Detected”. Reference intervals are presented in **Suppl. File 1, Table S30**.

### Diversity and taxonomic analyses

Species-level taxonomic relative abundance data were used for diversity calculations, with a minimum relative abundance threshold of 0.05% applied. Alpha diversity was calculated using the Shannon index, and beta diversity was assessed using the Bray-Curtis dissimilarity metric (vegan package, version 2.7-1). Beta diversity was visualized with Non-metric Multidimensional Scaling (NMDS) plots.

### Sensitivity analysis excluding individuals with IBD

To assess the robustness of observed associations and to evaluate whether results were driven by individuals with severe inflammatory disease, a sensitivity analysis was performed excluding participants with a reported diagnosis of IBD. All alpha and beta diversity analyses were re-run in the No-IBD cohort using the same statistical methods, group definitions, and multiple-testing correction procedures as in the primary analysis.

Effect sizes, as measured by Glass’ rank biserial correlation for alpha diversity and R^2^ from PERMANOVA for beta diversity, from the full cohort and No-IBD cohort were compared to evaluate consistency in the direction and magnitude of associations. Differences observed in the sensitivity analysis were interpreted in the context of baseline diversity differences between cohorts.

### Biomarker-normal healthy sensitivity analysis

Because the healthy reference group was based on self-reported data, we performed a sensitivity analysis using a stool chemistry-normal reference group. Self-reported healthy participants were classified as stool chemistry-normal if calprotectin was ≤ 80 µg/g, lactoferrin ≤

7.3 µg/g, and occult blood was negative (**Suppl. File 1, Table S30**); those outside the reference range for any of these three metrics were classified as stool chemistry-discordant. We quantified the discordance rate, compared Shannon diversity between stool chemistry-normal and stool chemistry-discordant participants using the Wilcoxon rank-sum test (with Glass’ rank-biserial effect size), and repeated the covariate-adjusted healthy-versus-condition analyses using each reference group. Alpha diversity was analyzed using linear models (Shannon ∼ group + age + BMI + sex), and beta diversity using PERMANOVA (adonis2; Bray–Curtis ∼ group + age + BMI + sex, by = “margin”), reporting the unique variance explained (R²) by group.

### Sex-stratified sensitivity analysis

To evaluate whether the female predominance of the cohort influenced the observed associations, we repeated the fully adjusted multivariable analyses separately in females (n = 668) and males (n = 209). Models included all covariates described above, with the exception of sex, which was omitted because analyses were stratified by sex. Four representative stool chemistry metrics (butyrate percentage, sIgA, stool pH, and calprotectin) were evaluated. Adjusted PERMANOVA *R*² values were compared between sexes, and the direction of association was assessed for taxa reaching q < 0.25 in both strata.

### Occult blood stratification

To evaluate whether occult blood positivity, including among self-reported healthy participants, could reflect menstrual contamination, we stratified occult blood positivity by sex and, within the self-reported healthy subset, by sex and age. Menstrual and menopausal status were available for only a small subset of female participants and were therefore not included in the analyses. To characterize microbiome differences associated with occult blood positivity, we compared Shannon alpha diversity (Wilcoxon rank-sum test) and Bray–Curtis community composition (PERMANOVA) between occult blood-positive and occult blood-negative participants in the overall cohort, within each sex, and within the self-reported healthy subset.

### Stool chemistry associations and network analysis

Significant associations between stool chemistry metrics and microbial features were identified using MaAsLin2^40^, with CLR transformed relative abundance data and the default parameters. Models were adjusted for age, sex, and BMI. Significant associations with FDR-adjusted q-values < 0.05 were used to construct the bipartite network, with nodes representing microbial features and stool chemistry covariates. To convert negative association coefficients into positive edges, stool chemistry nodes were split into “increased” and “decreased” categories (e.g., increased versus decreased butyrate). To focus on key features, the network was subset to include only nodes linked to increased or decreased levels of butyrate, calprotectin, or lactoferrin. Microbial features were visualized separately at different taxonomic levels to improve clarity.

To evaluate the robustness of stool chemistry-microbial associations to progressively more comprehensive confounder adjustment, we performed a nested series of MaAsLin2 analyses (using the same parameters as above) for all stool chemistry metrics at the species and genus levels. Models were fit sequentially as follows: M0 (age only), M1 (+ sex, BMI, and race/ethnicity), M2 (+ estimated dietary fiber intake), M3 (+ smoking and alcohol use), M4 (+ probiotic use), M5 (+ medication use), M6 (+ stool consistency and bowel movement frequency), and M7 (+ sequencing batch and sequencing depth; full model). At each step, we recorded the number of microbial features remaining significant at the MaAsLin2 default threshold (FDR-adjusted *q* < 0.25) and confirmed results using a more stringent threshold (*q* < 0.05). The direction of each association was tracked across all adjustment stages. The identical nested modeling framework was applied to both targeted assembly (PLASS) and untargeted k-mer functional profiles (funprofiler) to evaluate whether robustness to confounder adjustment generalized beyond taxonomic features.

### Machine learning classification of stool chemistry metric status

We developed an early-fusion elastic-net framework to predict stool chemistry metric status from whole-metagenome shotgun sequencing data. Each stool chemistry metric was modeled as two binary tail-classification tasks, a below-range task (Below range vs. Within/Above range) and an above-range task (Above range vs. Within/Below range); occult blood was modeled as a binary presence/absence outcome. This yielded 19 evaluable tasks across the chemistry panel. Four headline targets (occult blood, high calprotectin, low butyrate percentage, high acetate percentage) are featured in the main text, with the complete panel reported in **Suppl. File 1, Table S25**.

#### Feature views and early fusion

Three microbiome feature sets (“views”) were used: species-, genus-, and family-level taxonomic relative abundances. Functional features — targeted functional-gene abundances (KEGG Orthology and CAZy families) and KO modules quantified with funprofiler — were additionally evaluated in the model comparison but were not retained in the headline model. Taxonomic features were filtered using a minimum relative abundance threshold of 0.05% and a prevalence threshold of 5% across samples; functional features were filtered at a minimum prevalence of 5%. Taxonomic features were transformed using the centered log-ratio (CLR) transformation to account for compositionality: abundances were floored at 5 × 10⁻⁴, with zero values replaced by a constant pseudocount of 2.5 × 10⁻⁴ (half the floor) prior to the log-ratio. Feature selection was performed independently within each training partition.

#### Model comparison

Model choice followed a comparison of the elastic-net logistic regression (glmnet package, α = 0.5) on the CLR-transformed taxonomic features against random forest and gradient-boosted trees (LightGBM) compared taxonomy alone against taxonomy combined with the functional views in a single early-fusion matrix; and compared early fusion against a late-fusion stacked ensemble (LightGBM base learners combined by an elastic-net meta-learner). Configurations were ranked by nested-CV AUPRC. The elastic-net matched the best-performing alternatives on discrimination (panel-wide mean ΔAUPRC ≈ 0 versus LightGBM) while providing interpretable coefficients and a single-stage fit, and was retained as the reference model.

#### Cross-validation and threshold optimization

Performance was estimated using grouped, class-stratified nested cross-validation. Outer folds were grouped on purchasing account so that samples from the same account or household never spanned the train/test split. Inner folds were used solely for hyperparameter and threshold selection. Classification thresholds were selected on inner-fold predictions, fixed, and applied once to the corresponding outer test fold; no threshold selection was informed by test-fold outcomes.

#### Performance evaluation

Model performance was assessed using multiple, complementary metrics. AUPRC (PRROC package) was the primary selection and reporting metric, reported alongside AUROC, sensitivity, specificity, positive and negative predictive value, Cohen’s κ, balanced accuracy, and decision-curve net benefit. For clinical interpretation, each model was additionally evaluated at a high-sensitivity operating point (target sensitivity ≈ 0.90), with the resulting specificity, predictive values, and net benefit reported.

#### External validation

The elastic-net was retrained on the full discovery cohort, and the fixed transform, coefficients, and high-sensitivity threshold were applied without refitting to a subject-disjoint external cohort of 500 kits (no shared accounts; sampled after the discovery-cohort freeze). Internal-versus-external AUROC and AUPRC, and transfer of the operating point, are reported.

#### Alternative models

Two alternatives to binary tail classification were evaluated. A three-class model (Below/Within/Above range) was assessed by Cohen’s κ. An elastic-net cumulative-logit (proportional-odds) model was compared against the paired dichotomized tail classifiers by AUPRC and Kendall’s τ. To characterize what the microbiome can predict quantitatively, raw biomarker concentrations were regressed on the CLR(taxa + function) features using elastic-net, reporting R² and Spearman’s ρ. All models were fitted in R (version 4.5.1).

### Statistical analyses

All statistical analyses were performed in R (version 4.5.1). Summary statistics for the cohort were generated using the gtsummary package (version 2.3.0). Differences in alpha diversity between stool chemistry classification groups (“Below,” “Within Range,” “Above”) were assessed using the Wilcoxon Rank-Sum test, with p-values adjusted for multiple comparisons using the Benjamini-Hochberg (FDR) method.

To characterize the relationships among the stool-chemistry metrics themselves, we computed pairwise Spearman rank correlations between all continuous metrics (Benjamini–Hochberg-adjusted) and, for the binarized out-of-range calls, pairwise Fisher’s-exact tests of co-abnormality (odds ratios, BH-adjusted).

To evaluate whether associations between stool chemistry metric and microbiome composition persisted after adjustment for potential confounders, we performed multivariable variance partitioning using distance-based redundancy analysis (dbRDA, vegan). For each stool chemistry metric, the unique (partial) variance explained (partial R²) was estimated while adjusting for demographic variables (age, sex, BMI, and race/ethnicity), dietary fiber intake (estimated using self-reported weekly plant-based food servings), probiotic use, medication use (NSAIDs, PPIs, and laxatives), stool consistency and transit time (Bristol Stool Form Scale and bowel frequency per day, both continuous), sequencing batch (measured as continuous months sample was sequenced), and sequencing depth (log-transformed total sequencing reads). Missing medication data were modeled as a separate category.

Statistical significance was assessed using 499 permutations, and p-values were adjusted using the Benjamini–Hochberg method across stool chemistry metrics within each microbiome feature set (taxonomy, targeted functional genes, KO modules). Unique partial R² values were similarly calculated for each confounder while conditioning on all remaining variables in the full model. For comparison with the primary analyses, marginal (unadjusted) PERMANOVA models were also performed using vegan::adonis2 with 999 permutations, controlling only for age and BMI.

Homogeneity of group dispersions was assessed using the vegan::betadisper function. Pairwise Wilcoxon Rank-Sum tests were applied to compare taxon abundances across biomarker groups, with p-values corrected for multiple testing using the FDR method. For selected figures, extreme outlier values were excluded from visualization only to improve interpretability; no data points were excluded from statistical analyses.

## Data availability statement

Deidentified individual participant data and microbial reads generated and analyzed for this manuscript are available in the National Center for Biotechnology Information (NCBI) repository under bioproject accession PRJNA1428157.

## Author contributions

**Tim Straub**: Conceptualization, formal analysis, methodology, visualization, writing - review & editing. **Pamela A. Nieto**: Conceptualization, writing - original draft, writing - review & editing, visualization. **Julian Trachsel**: Conceptualization, formal analysis, methodology, visualization, writing - review & editing. **Hannah Sturgeon**: writing - original draft, writing - review & editing. **Emeran Mayer, Ruben A.T. Mars, Star Edwards & Cheryl S. Hoy:** Writing - review & editing. **Kimberley V. Sukhum**: Conceptualization, supervision, writing - review & editing.

