## Supplemental file 2 for "Integrating stool chemistry measurements with metagenomics in adults reveals distinct gut health profiles"

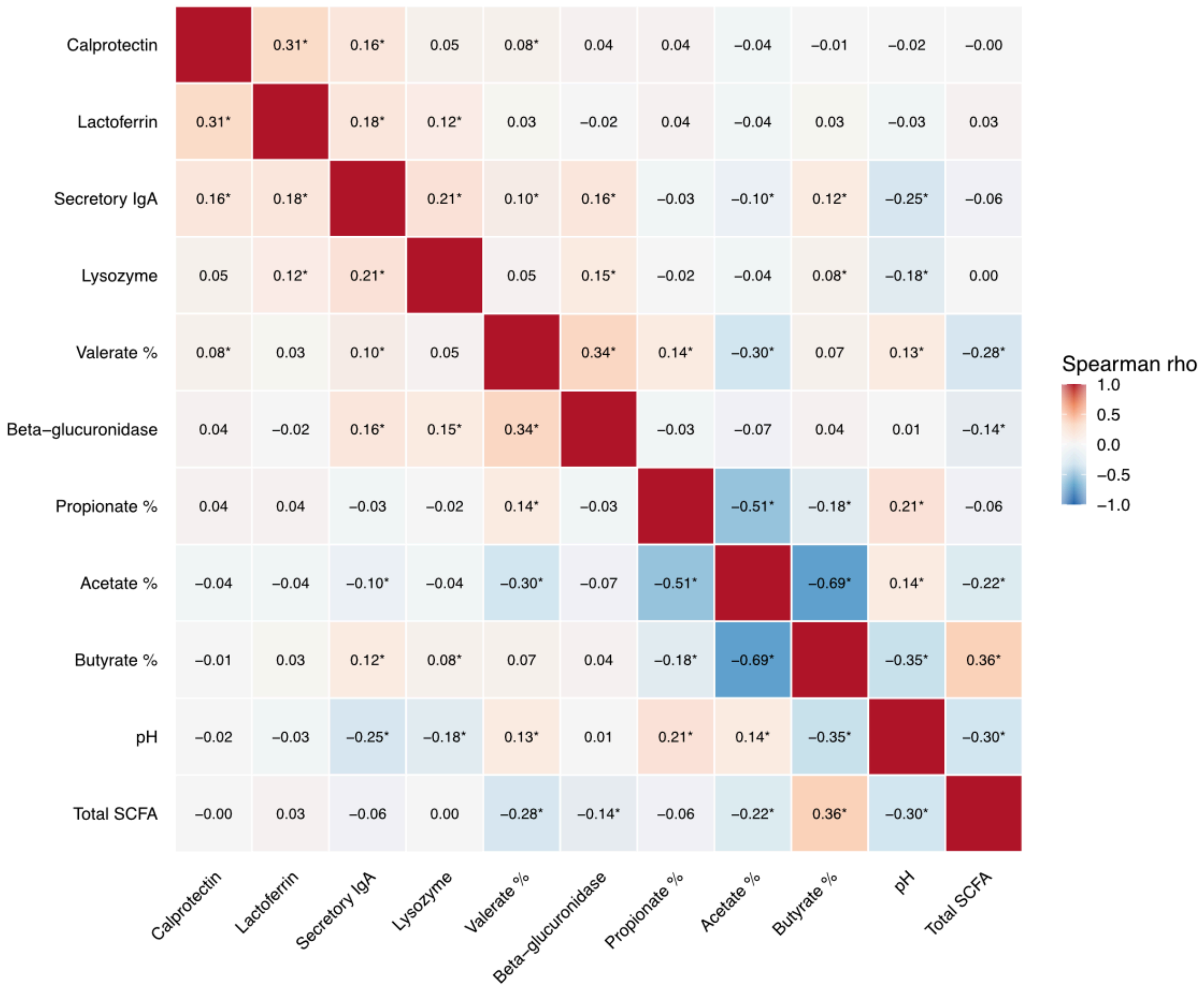


**Figure S1: Pairwise correlations among continuous stool chemistry metrics.** Spearman rank correlation coefficients between continuous stool chemistry metrics are displayed as a heatmap. The color gradient represents the direction and strength of association, with blue indicating negative correlations and red indicating positive correlations. Asterisks indicate statistically significant correlations after Benjamini–Hochberg FDR correction ($q < 0.05$).

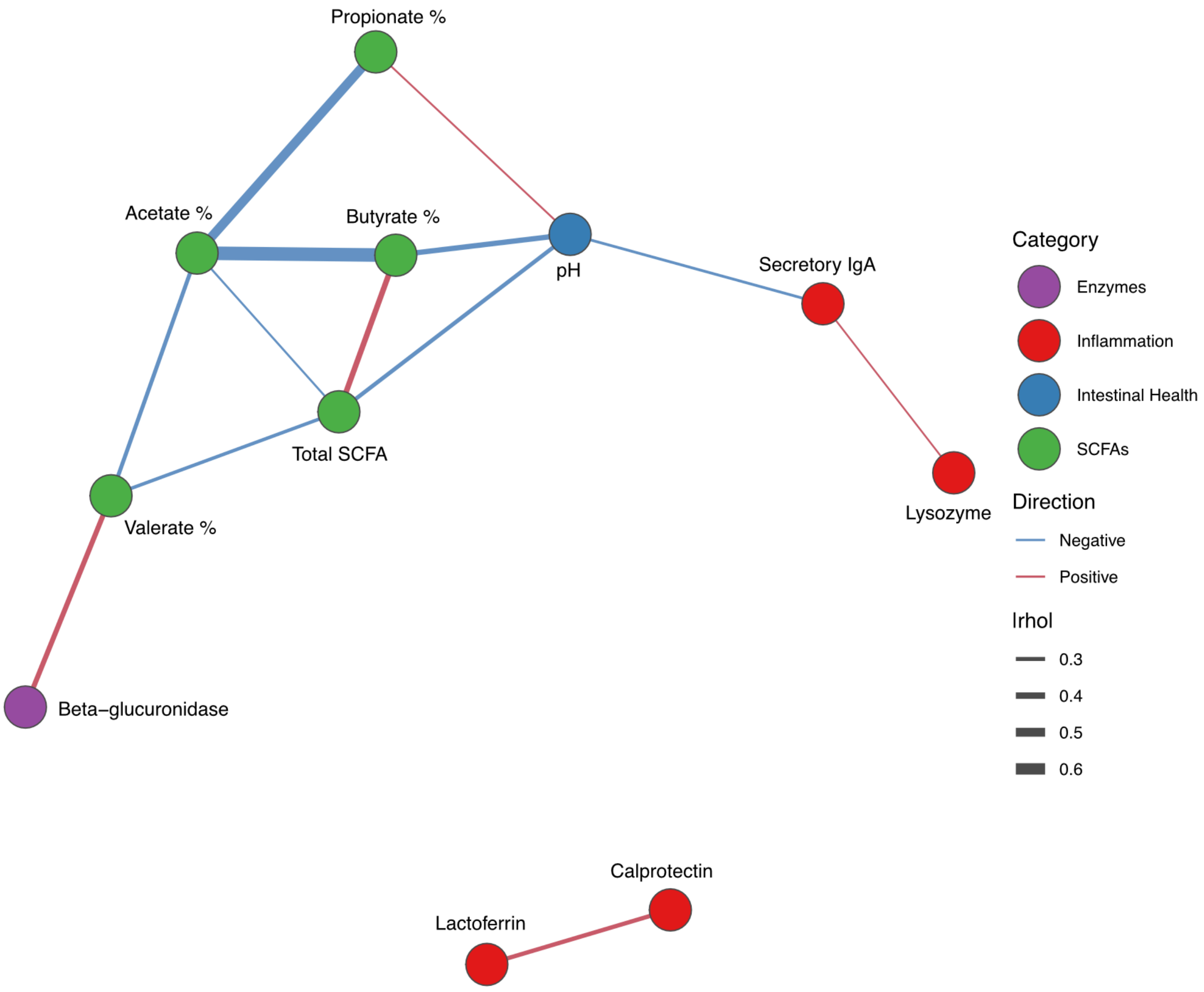


**Figure S2: Correlation network of stool chemistry metrics**. Network visualization of significant pairwise correlations among stool chemistry metrics. Nodes represent individual metrics and are colored according to category. Edges represent correlations meeting significance and effect-size thresholds (Benjamini–Hochberg FDR–adjusted $q < 0.05$ and $|\rho| \geq 0.2$). Edge color indicates the direction of association, with red representing positive correlations and blue representing negative correlations. Edge width is proportional to the absolute magnitude of the Spearman correlation coefficient ($|\rho|$), with thinner lines indicating weaker and thicker lines indicating stronger associations.

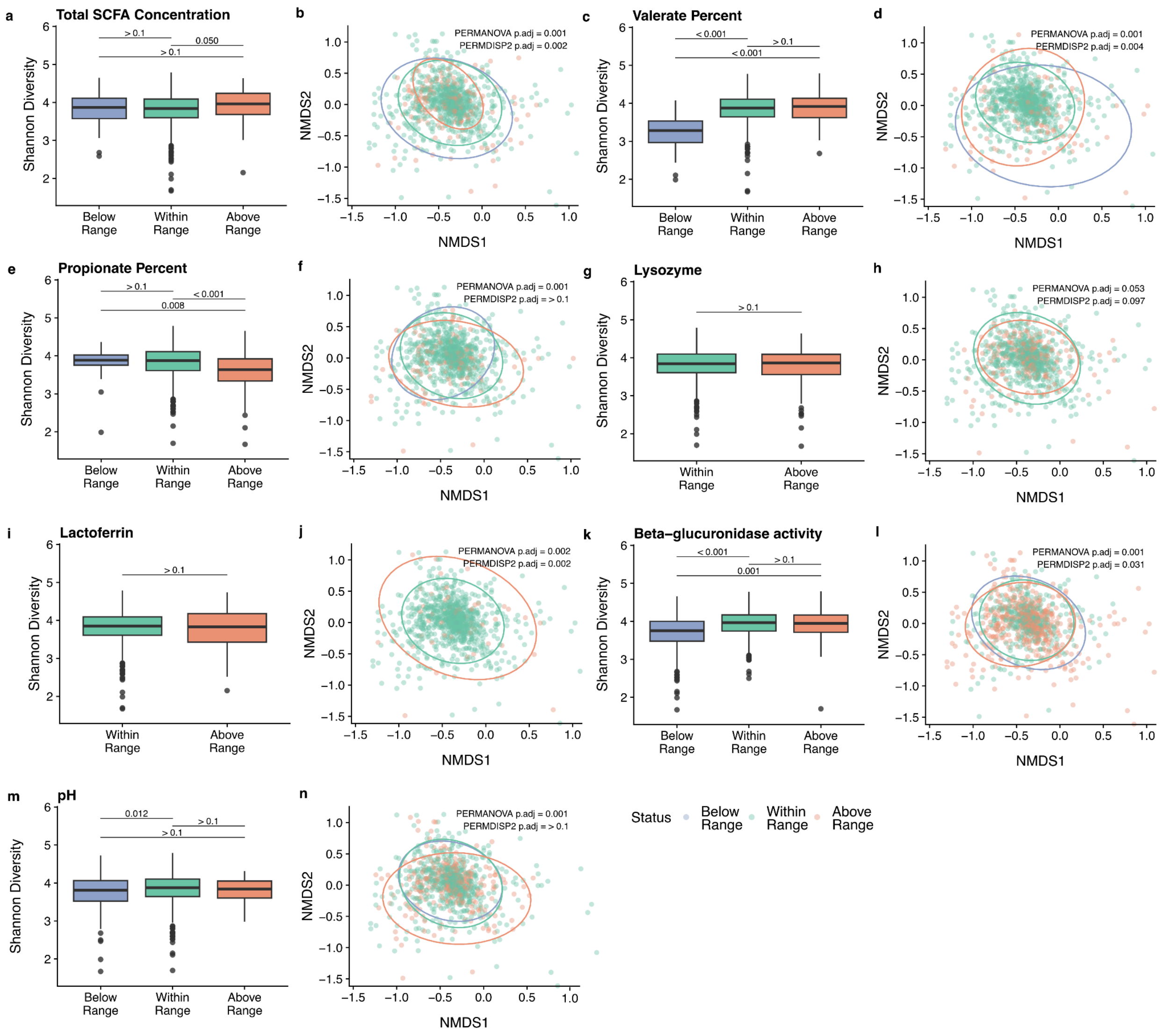


**Figure S3: Gut microbiome diversity and composition across different stool chemistry metrics.** Shannon diversity (left panels) and non-metric multidimensional scaling (NMDS; right panels) plots showing gut microbiome variation according to stool chemistry metrics, including **(a, b)** total SCFA concentration, **(c, d)** valerate percent, **(e, f)** propionate percent, **(g, h)** lysozyme, **(i, j)** lactoferrin, **(k, l)** beta-glucuronidase activity, and **(m, n)** pH. Samples were grouped as below range (blue), within range (green), or above range (orange) based on reference ranges for each chemistry metric. Boxplots display Shannon diversity indices, with statistical differences assessed using the Wilcoxon rank-sum test and Benjamini–Hochberg–adjusted *p*-values indicated above group comparisons. NMDS ordinations are based on Bray–Curtis dissimilarities, with ellipses indicating 95% confidence intervals. Statistical significance in community composition was tested using PERMANOVA, and differences in dispersion were evaluated using PERMDISP2. Two outlier values were omitted from these plots to improve visualization; all statistical analyses were performed on the full dataset.

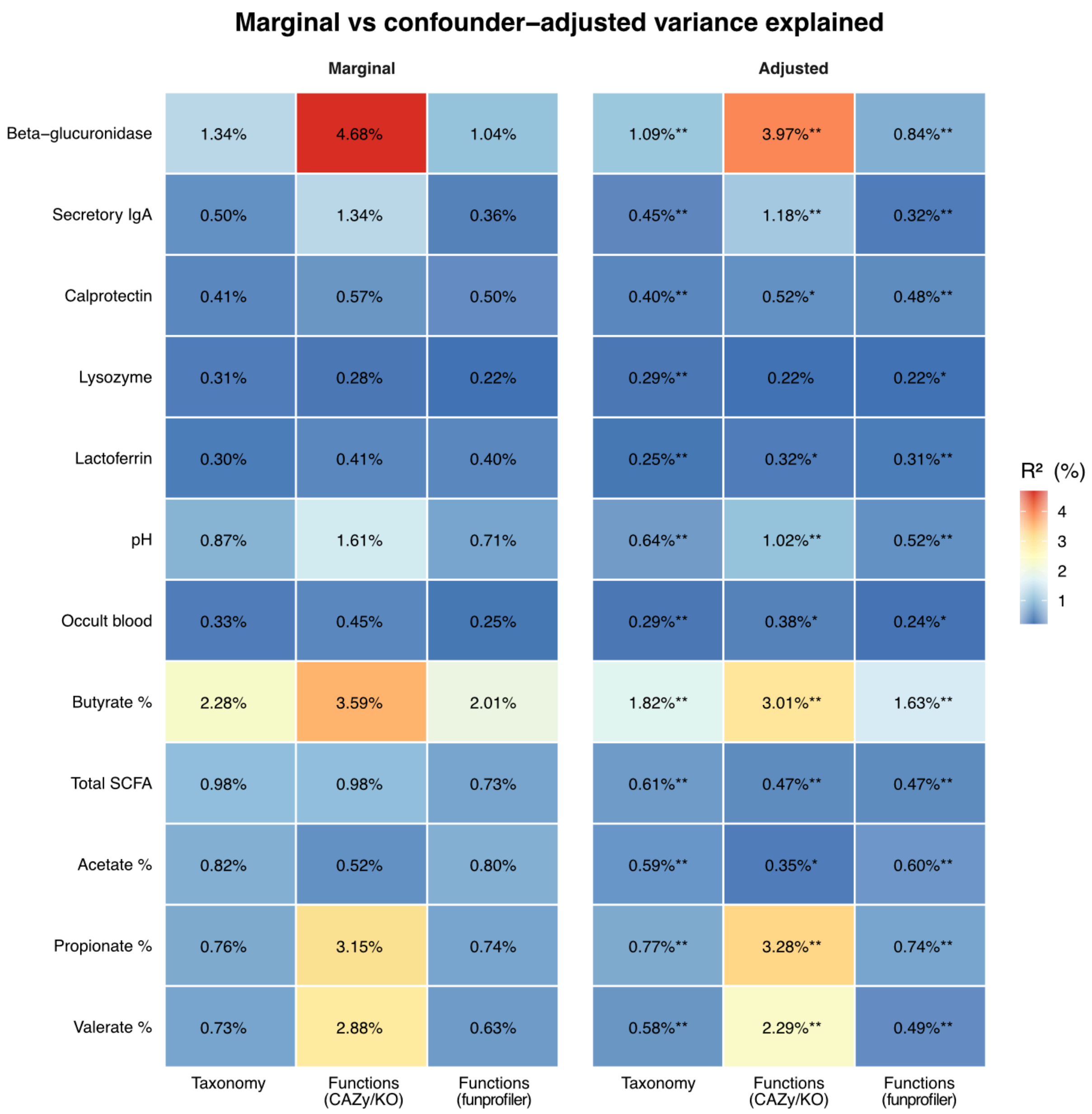


**Figure S4: Variance explained across taxonomic and functional profiles.** Heatmap showing the marginal and confounder-adjusted proportion of variance explained ($R^2$) by stool chemistry metric across three data types: taxonomic composition, targeted functional profiles (CAZy/KO), and inferred functional profiles (FunProfiler). Values represent the percentage of variation explained by each factor, with warmer colors indicating higher explanatory power. Asterisks denote statistical significance (* $p < 0.05$; ** $p < 0.01$).

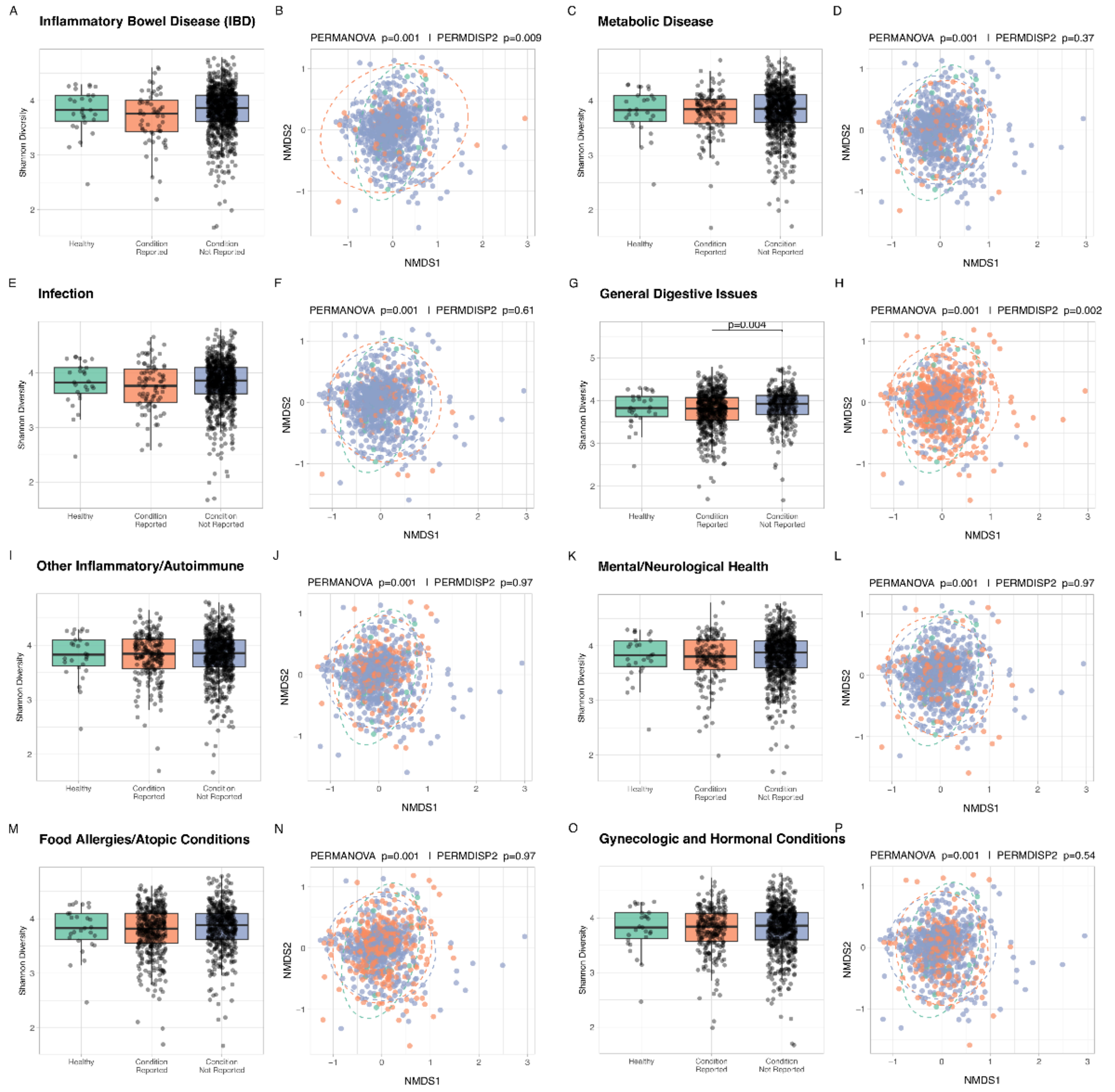

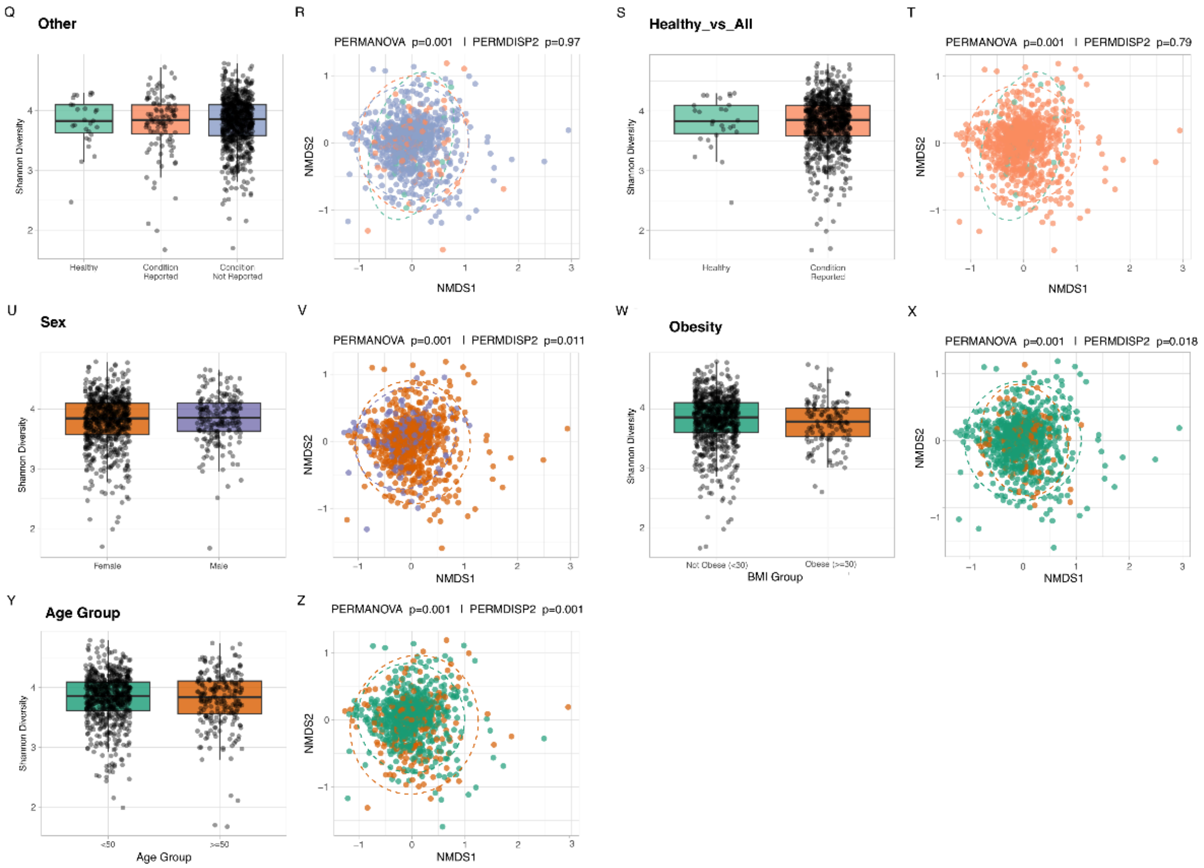


**Figure S5: Gut microbiome diversity and composition across different health conditions.** Shannon diversity (left panels) and non-metric multidimensional scaling (NMDS; right panels) plots showing gut microbiome variation according to self-reported health conditions, including **(a, b)** Inflammatory bowel disease, **(c, d)** metabolic disease, **(e, f)** infection, **(g, h)** general digestive issues, **(i, j)** other inflammatory/autoimmune conditions, **(k, l)** mental/neurological conditions **(m, n)** food allergies/atopic conditions, **(o, p)** gynecologic and hormonal conditions, **(q, r)** other conditions. Diversity and NMDS plots are also shown for samples grouped by **(u, v)** sex, **(w, x)** BMI, and **(y, z)** age.

For conditions, samples were grouped as healthy (green), condition reported (orange), or condition nor reported (blue). Boxplots display Shannon diversity indices, with statistical differences assessed using the Wilcoxon rank-sum test and Benjamini–Hochberg–adjusted *p*-values indicated above group comparisons when significant. NMDS ordinations are based on Bray–Curtis dissimilarities, with ellipses indicating 95% confidence intervals. Statistical significance in community composition was tested using PERMANOVA, and differences in dispersion were evaluated using PERMDISP2.

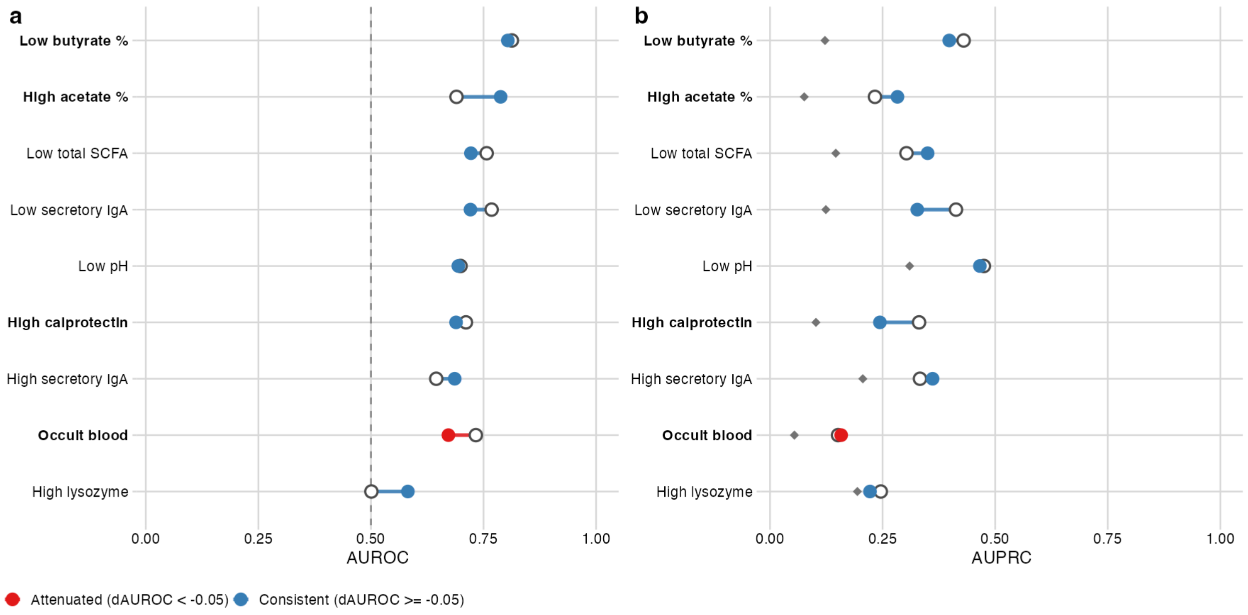


**Figure S6: Internal-to-external classifier discrimination across the external-validation panel** (**Suppl. File 1, Table S27**). **(a)** AUROC and **(b)** AUPRC for each target; open circles = internal estimate (877-kit discovery cohort, pooled out-of-fold predictions), filled circles = external estimate (held-out 500-kit validation cohort), connected by a line segment. Segment/filled-circle colour indicates whether external AUROC held within 0.05 of internal (blue, consistent) or dropped by more (red, attenuated). Grey diamonds in (b) mark each target's external-cohort prevalence, the chance baseline for AUPRC. Dashed vertical line in (a) marks chance (AUROC = 0.5). Bold row labels denote the four metrics reported in **Figure 6**.
